# Multicellular Spatial Programs Define the Histopathological Architecture of Meningioma

**DOI:** 10.64898/2026.06.16.732172

**Authors:** Danielle F. Miyagishima, Arman Afrasiyabi, Declan McGuone, E. Zeynep Erson-Omay, Kanat Yalcin, Yutaka Takeo, Phan Q. Duy, Batur Gultekin, Jacky Yeung, A. Gulhan Ercan-Sencicek, Octavian Henegariu, Mark W. Youngblood, Ketu Mishra-Gorur, Katsuhito Yasuno, Guilin Wang, Nenad Sestan, Roel G.W. Verhaak, Jennifer Moliterno, Tanyeri Barak, Murat Gunel

## Abstract

Tumors are often described by cell types or gradients, but the organizing units of tumor tissue remain unclear. Using meningiomas, which show marked morphologic diversity despite constrained and recurrent genetics, we identify reproducible multicellular spatial molecular programs (SMPs) characterized by recurring cell-type mixtures that combine in different proportions across tumors. We built a multi-omic atlas of 147 human meningiomas (1.6 million cells/spots), integrating scRNA-seq, Visium, CosMx-RNA, and CosMx-Protein. Across platforms, SMPs mapped onto canonical whorl–lobule architecture and defined a structured ecological landscape linking hypoxic, immune-evasive states to vascularized, matrix-remodeling, and mineralization-rich states. For translation, we developed MeningNet, a hybrid ConvNeXt–Vision Transformer that infers SMPs directly from hematoxylin-and-eosin sections. MeningNet generalized across an internal replication cohort and 465 external whole-slide images, recovering cross-platform inference and showing significant association with CNS-WHO grade. These findings establish meningioma architecture as a reproducible histopathological framework for inferring spatial molecular state from routine pathology.

## Introduction

Tumor progression reflects not only the accumulation of somatic mutations but also the dynamic interplay between mutation-bearing cells and the spatially organized microenvironments in which they reside.^1,2^ Spatial gradients of hypoxia, nutrient availability, and immune pressure impose selective forces that shape clonal evolution across space and time.^3–6^ Spatially resolved transcriptomic and proteomic analyses further demonstrate that genomic programs and ecosystem architecture are interdependent components of tumor behavior.^7–9^ However, in many tumor types, extensive mutational complexity obscures the relative contribution of microenvironmental architecture to phenotypic heterogeneity, limiting our ability to disentangle genetic drivers from ecological context.

Meningiomas provide a uniquely tractable system for examining this relationship. As the most common primary intracranial tumors,^10^ they exhibit low mutational burden and recurrent, largely mutually exclusive somatic driver alterations, including *NF2*, *TRAF7*, *KLF4*, *AKT1*, *SMO*, and *POLR2A*, that define major molecular groups.^11–16^ At the same time, they preserve stereotyped microanatomical structures, including whorls and lobules, across grades and histologic variants.^17^ This combination of constrained genetics and conserved tissue morphology creates an opportunity to interrogate how genomic background, microenvironmental composition, and spatial architecture interact to generate clinically meaningful heterogeneity.

While somatic mutation status associates with histology, anatomical location, and grade, the most common mutation subgroup, *NF2*-mutant/Chr22q loss meningiomas, exhibits divergent outcomes based on location and differences in immune infiltrate.^18^ Single-cell RNA-seq studies in meningioma have begun to map this cellular landscape, defining stromal and immune compartments^19,20^ and, more recently, methylation- and microenvironment-resolved cell-state taxonomies.^21,22^ These studies establish that microenvironmental composition shapes molecular classification and challenge rigid categorical models of tumor identity. However, because dissociated single-cell approaches do not preserve native tissue architecture, the spatial organization of these states within intact microanatomy, and their relationship to canonical histologic structures, remain incompletely defined.

Together, these findings raise a central question: do tumor-level molecular continua reflect unstructured variation, or do they arise from the differential representation of discrete, spatially organized tissue states embedded within canonical microanatomy? Pan-cancer single-cell analyses have identified conserved malignant programs and emphasized both discrete and continuum-like tumor cell states.^23,24^ Yet the architectural question, namely what reproducible multicellular units organize tumor tissue, has not been directly addressed in meningioma or in solid tumors more broadly.

Here, we construct a spatially resolved, multi-omic atlas of 147 human meningiomas spanning all CNS WHO grades, major histologies, and genomic backgrounds. By integrating single-cell transcriptomics (220,043 cells from 20 tumors and 3 dura from 18 patients), spatial transcriptomics (10x Visium; 56,020 spots from 17 sections from 10 patients), single-cell spatial transcriptomics (CosMx RNA; 391,051 cells from 77 patients), and spatial proteomics (CosMx Protein; 993,675 cells from 119 patients) with matched histology, we define reproducible multicellular spatial molecular programs (SMPs) that organize tumor tissue architecture across patients and platforms. These programs define stable architectural units that integrate cellular composition with microanatomical organization. Among these, we identify a whorl–lobule–associated state linking canonical histologic structures to coordinated epithelial–mesenchymal transition (EMT), hypoxia, and immune-associated programs.

A key limitation of spatial transcriptomic approaches is their limited scalability and lack of availability in routine clinical practice. To overcome this, we developed an interpretable deep learning framework that infers spatial molecular programs directly from standard H&E sections, enabling recovery of spatial molecular architecture without molecular profiling. We demonstrate robust generalization across an internal replication cohort of 128 tumors and an external cohort of 465 whole-slide images from multiple imaging platforms, enabling prediction of tumor grade and recurrence without retraining.

Together, these findings establish spatial molecular programs as a reproducible framework linking morphology, molecular state, and clinical behavior, and demonstrate that tumor ecosystem architecture can be decoded directly from routine histopathology.

## RESULTS

### An integrated multi-omic atlas defines the cellular landscape of meningioma

To assess the compositional heterogeneity of meningiomas, we created a multi-omic atlas comprising 147 tumors from 141 individuals (Supplementary File 1). This included single-cell transcriptomic sequencing (scRNA-seq) from 220,043 cells, whole-transcriptome spatial sequencing (10x Visium) from 56,020 spots from 17 tumor sections, single-cell resolution spatial transcriptomic sequencing (NanoString CosMx RNA) of 391,051 cells from 77 tumors, spatial proteomic profiling (NanoString CosMx protein) of 993,675 cells from 119 tumors (Figure 1 A).

**Figure 1.**
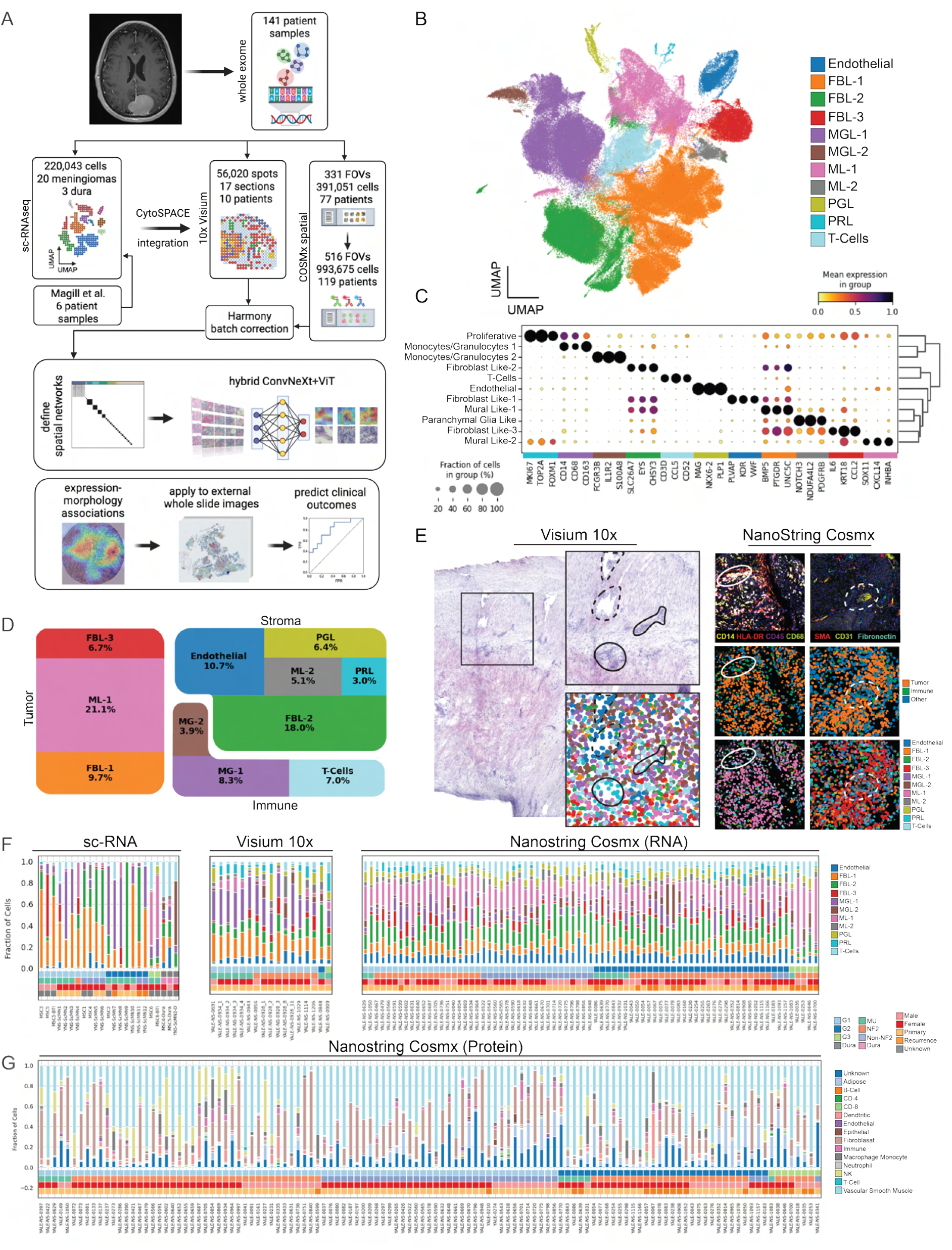
Spatially resolved multi-omic atlas of human meningioma reveals compositional heterogeneity. **A.** Overview of the study design and data integration workflow. Created in BioRender https://BioRender.com/od1nt6q. **B.** UMAP projection of scRNA-seq data showing 11 major cell populations, including fibroblast-like (FBL-1, FBL-2, FBL-3), mural-like (ML-1, ML-2), monocyte–granulocyte (MG-1, MG-2), endothelial, parenchymal glial-like (PGL), proliferative (PRL), and T cells. **C.** Dot plot showing expression of canonical marker genes across scRNA-defined cell populations, with dot size indicating the proportion of cells expressing each gene and color representing mean scaled expression. **D.** Voronoi diagram illustrating the relative proportions of tumor, immune, and stromal or other cell categories across the scRNA-seq dataset. **E.** Representative 10x Visium section showing spatial localization of scRNA-defined cell types following CytoSPACE integration. Higher-magnification views highlight regions of vasculature (dotted outlines) and immune infiltration (solid outlines). Corresponding NanoString CosMx protein-based cell-type annotations are shown, with immune markers (CD45, HLA-DR, CD14, CD68) highlighting infiltrates and vascular or stromal markers (SMA, CD31, fibronectin) defining vasculature and fibroblast-rich regions. **F.** Bar plots showing cell-type composition across patients in the scRNA-seq, Visium, and CosMx RNA modalities. **G.** Cell-type distributions derived from NanoString CosMx protein annotations across patients, with associated clinical annotations including tumor grade, genomic subgroup, sex, and recurrence status.

#### Defining cell type populations present across meningiomas

Next, we performed scRNA-seq on 12 meningioma samples and one dura sample and integrated these data with 10 samples from a previously published dataset, comprising 8 tumors and 2 dura samples from 6 patients,^19^ for a total of 20 tumors and 3 dura samples from 18 individuals (Supplementary File 1). Unsupervised Leiden clustering followed by cluster merging based on transcriptional similarity (Pearson r > 0.5; Supplementary Figure S2, Supplementary File 2) identified 11 reproducible cell populations: fibroblast-like (FBL)−1, −2, and −3; mural-like (ML)-1 and −2; monocyte/granulocyte (MG)-1 and −2; endothelial; proliferative (PRL); parenchymal glial-like (PGL); and T-cell populations (Figure 1B). Marker genes and log_2_ fold expression changes were determined using EcoTyper,^25^ and cell population annotations were benchmarked against published meningioma single-cell taxonomies and cell annotation databases (Figure 1C, Supplementary File 3).^20,21,26^

To distinguish tumor from non-tumor populations, we inferred copy-number alterations from the scRNA-seq data using InferCNV, with dura-derived cells used as reference controls. Among the 20 scRNA-seq meningiomas, 11 harbored NF2 alteration and/or chromosome 22q loss; 7 of these had whole-exome sequencing available, and 5 of 7 showed recurrent chromosome 1p loss and additional less frequent chromosomal alterations (Supplementary File 1). InferCNV analysis identified consistent chromosome 22q loss in FBL-1, FBL-3, and ML-1 cells, with recurrent chromosome 1p loss in the same populations, supporting their assignment as tumor-cell populations (Supplementary Figure S3). In contrast, FBL-2, ML-2, MG-1, MG-2, endothelial, PGL, T-cell, and PRL populations lacked these broad arm-level alterations and were assigned as stromal, immune, endothelial, glial-associated, or cycling non-tumor populations according to their marker profiles and reference annotations.

FBL-1 and FBL-2 cells were enriched for genes previously reported in meningioma, including *LEPR* and *SLC4A4*, with FBL-1 further defined by higher expression of *PTGDR* and *OGN*, and FBL-2 defined by enrichment of ion transport genes including *SLC26A7* and *KCNMA1*.^20,21^ FBL-3 cells expressed inflammatory and chemotactic genes, including *CCL2*, *ICAM1*, and *IL6*, consistent with a cancer-associated fibroblast-like inflammatory state and overlapping with inflammatory fibroblast programs described in prior meningioma single-cell studies.^20,27^ ML-1 cells expressed *COL5A1* and *CXCL14*, whereas ML-2 cells were characterized by high expression of mural-cell markers including *RGS5* and *NOTCH3*. MG-1 cells exhibited canonical tumor-associated macrophage markers, including *CD163* and *MSR1*, whereas MG-2 cells expressed *FCGR3B*, *IL1R2*, and *S100A8*, consistent with an inflammatory monocyte/granulocyte-like state. Endothelial cells were annotated based on enriched expression of *KDR* and *PLVAP*. PGL cells were defined by high expression of glial-associated genes including *NKX6-2* and *PLP1*. T cells expressed canonical lymphoid markers including *CD8A*, *CCL5*, and *CD2*. PRL cells were distinguished by robust expression of cell-cycle genes including *MKI67*, *TOP2A*, and *FOXM1*, consistent with active proliferation.

To further characterize tumor cell presence, we assessed cell cycle activity using a standard G1/S/G2M gene signature.^28^ Cells were classified as cycling if they had a positive G2M or S score. Across all profiled cells within each grade class, the combined fraction of G2M- or S-phase cells increased with grade (G1 20.5%, G2 33.9%, G3 67.6%) (Supplementary Figure S4 A, Supplementary File 2). These fractions reflect the combined G2M + S population across all profiled cells per grade class, rather than the fraction within a defined cell state, and are consistent with prior reports of elevated proliferative programs in biologically aggressive meningiomas.^21^ This cohort-level estimate is therefore not directly comparable to per-state proliferative fractions reported in glioblastoma, where cycling activity was quantified within defined malignant cell states.^29^ In our cell-state-specific analysis, the Proliferative cell cluster had the highest proportion of cells assigned to G2M (81.2%) and S phase (18.8%) (Supplementary Figure S4 B). Notably, ML-1 cells also showed a high proportion of cycling cells, with 71.8% of cells in G2M or S phase (Supplementary Figure S4 B).

We used CellChat weighted directed network graphs to reconstruct intercellular signaling and identified that FBL-3, ML-1, and Proliferative cells were the major hubs of outgoing and incoming signals (Supplementary Figure S4 C-D).^30^ Among the top signaling pathways were TGF*β*, BMP, WNT, ncWNT, and EGF, which have been previously implicated in meningioma pathogenesis (Supplementary File 4).^31–34^ Taken together, the chromosomal aberrations, proliferative activity, and signaling centrality in FBL-1, FBL-3, and ML-1 cells support their designation as meningioma tumor cell populations (Figure 1 D).

Next, we identified transcriptional programs shared within and across tumor profiles by integrating the Visium and CosMx datasets. After preprocessing each modality independently, we restricted analyses to shared genes and performed batch correction using Harmony.^35^ Cross-platform mapping of scRNA-seq-defined cell types was achieved using CytoSPACE, enabling spatial resolution of cell-type abundance across platforms (Figure 1 E-G, Supplementary File 8).

#### Spatial molecular programs represent distinct multicellular units

To capture robust and recurring transcriptional patterns, we applied a previously validated framework for deriving meta-programs (MPs).^36^ Briefly, for each tumor, we generated gene sets using both non-negative matrix factorization (NMF, k = 2-11) and Leiden clustering (resolution = 0.5) (Supplementary File 5). This yielded 2,372 gene programs across all 94 spatial RNA-sequenced tumors (17 Visium sections, 77 CosMx tumors), which were compared using Jaccard similarity. Consensus MPs were defined by identifying gene sets that were recurrent across tumors, resulting in 18 MPs, each comprising 50 genes (Figure 2A, Supplementary File 5). Consistent with prior observations of extensive inter-individual heterogeneity in meningioma at the single-cell level,^37^ approximately 33% of programs were not consistently assigned across tumors, limiting the generalizability of discrete MPs alone.

**Figure 2.**
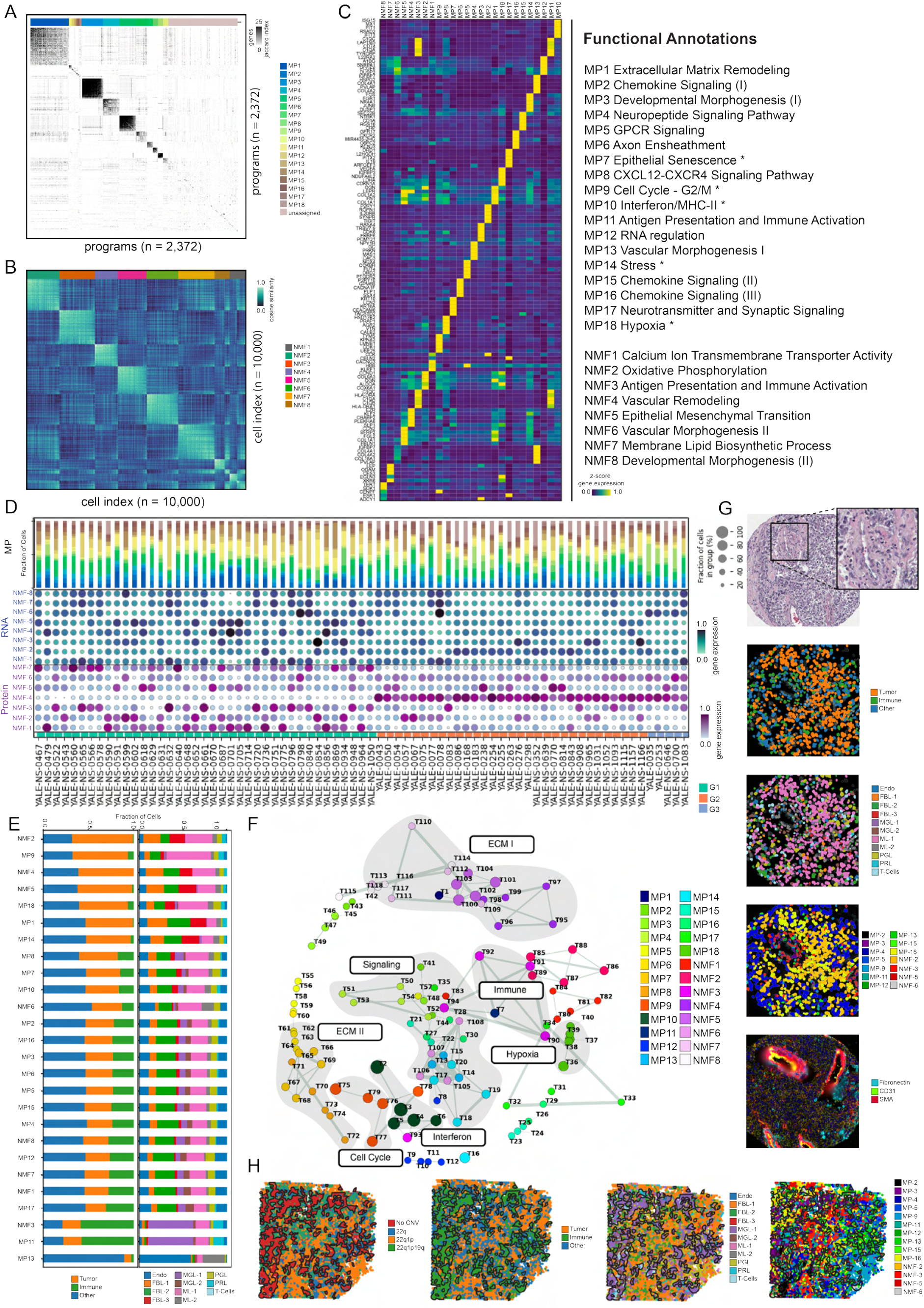
Spatial molecular programs define recurrent, multicellular tissue states that extend beyond single-cell classifications. **A.** Jaccard similarity matrix of gene programs derived from per-sample Leiden clustering and NMF across spatial transcriptomic sections (n = 94), used to identify recurrent meta-programs (MPs). **B.** Cosine similarity matrix of RNA-based NMF expression profiles across combined Visium and NanoString datasets (subsampled to 10,000 cells for visualization), ordered by dominant NMF component, illustrating robust component structure across individual cells. **C.** Heatmap of the top five genes per MP and RNA-NMF component (z-scored by expression), with functional annotations shown at right. Asterisks denote programs overlapping previously reported pan-tumor meta-programs. **D.** Program usage across tumor cells. Columns correspond to individual tumors. Top: MP assignments; middle: RNA-based NMF component assignments; bottom: protein-based NMF component assignments, highlighting concordance and divergence across molecular modalities. **E.** SMPs ranked by fraction of tumor cells (left) and corresponding cellular composition of each SMP (right). **F.** Network representation of enriched gene sets from top-ranked MPs and RNA-NMF components. Nodes represent gene sets (labeled by shorthand identifiers; see Supplementary File 5), with size proportional to enrichment significance (–log_10_p) and color indicating the originating SMP. Connecting edges denote Jaccard similarity between gene sets, with thickness proportional to gene overlap. Asterisks denote correspondence to previously described pan-tumor programs. **G.** Representative H&E image centered on a vascular structure with aligned spatial annotations, illustrating how cell-type labels, SMP assignments, and protein expression patterns collectively represent tissue architecture. **H.** Yale-NS-0856 sample with (from left) spatially-derived CNV identified by inferCNV, categories derived from scRNA-based inferCNV categories, cell-type classification using CytoSPACE integration, and joint SMP categorization (right).

To overcome this limitation, we applied a continuous NMF decomposition across all spatial RNA samples (Figure 2 B). Most NMF components were functionally distinct from the consensus MPs, with only 2 of 18 MPs showing a Pearson correlation r > 0.5 with any NMF component (Supplementary Figure S5, Supplementary File 5). Analysis of the ontological terms associated with the top 50 genes revealed enrichment for distinct biological functions, supporting that SMPs cannot be reduced to cell-type composition or to a single program family, but rather emerge as multicellular units of tissue organization (Figure 2 C, Supplementary Figure S6). We refer to these combined representations as spatial molecular programs (SMPs), which integrate discrete recurrent gene programs with continuous spatial gradients, enabling coordinated transcriptional states to be shared across tumors yet variably arranged within tissue. Because CosMx profiling provides matched spatial protein measurements from adjacent tissue sections, we assessed whether analogous spatial programs could be independently recovered at the protein level. Due to the difference between the profiled proteins (n = 64) and RNA transcripts (n = 6,000), we performed an independent NMF decomposition of the protein expression matrix, identifying seven protein-based components (Supplementary Figure S8). These included immune, tumor, and stromal-associated programs (Figure 2 D).

When examined on a per-sample basis, we observed that the SMPs share characteristic compositions across tumors, with most components present to varying degrees across all samples (Figure 2 D). Furthermore, SMPs formed multicellular units of heterogeneous cellular composition (Figure 2 E, Supplementary File 5). No SMP corresponded to a single scRNA-defined cell type; instead, each program comprised characteristic mixtures of tumor, immune, and stromal populations, indicating that SMPs in meningioma capture higher-order tissue states rather than expanded cellular classes.

To better understand the relationship between SMPs, we constructed a network of gene sets enriched in the top programs using Jaccard similarity based on shared gene content (Figure 2F). Network clustering revealed major functional modules corresponding to immune signaling, extracellular matrix (ECM) remodeling, angiogenesis, cellular stress responses, and cell cycle regulation (Figure 2 F). We then used a hypergeometric test to compare the previously published pan-tumor MPs (not including meningiomas)^36^ and observed significant overlap with several previously defined pan-tumor programs, including MP10 (interferon/MHC-II; Jaccard = 0.23, adjusted p = 3.93 × 10^-37^), MP14 (stress; Jaccard = 0.22, adjusted p = 7.26 × 10^-35^), MP9 (cell cycle/G2M; Jaccard = 0.19, adjusted p = 6.15 × 10^-21^), MP18 (hypoxia; Jaccard = 0.17, adjusted p = 9.11 × 10^-28^), and MP7 (epithelial senescence; Jaccard = 0.12, adjusted p = 6.9 × 10^-19^) (Supplementary File 5). Two additional MPs (MP12 (RNA regulation) and MP1 (extracellular matrix remodeling)) overlapped with EMT-associated pan-tumor programs. Together, these analyses demonstrate that spatial molecular programs define biologically meaningful multicellular states that integrate and extend single-cell classifications by capturing coordinated spatial organization within meningioma tissue (Figure 2 G, H).

#### Spatial molecular programs organize tissue architecture and associate with aggressive tumor features

To determine how SMPs relate to histologic subtype and aggressive clinical behavior, we quantified program enrichment across meningioma histologies, CNS WHO grade, and recurrence status (Figure 3, Supplementary File 8). We also assessed for differences in spatial configuration (Figure 3 A). For each tumor, we calculated the fraction of cells assigned to each program based on the dominant component per cell and compared groups using Wilcoxon rank-sum testing (Figure 3 B, D, and Supplementary File 8). Meningothelial, transitional, and atypical tumors were analyzed in detail, as these histologies are representative of high- and low-grade and have >10 tumors in our cohort.

**Figure 3.**
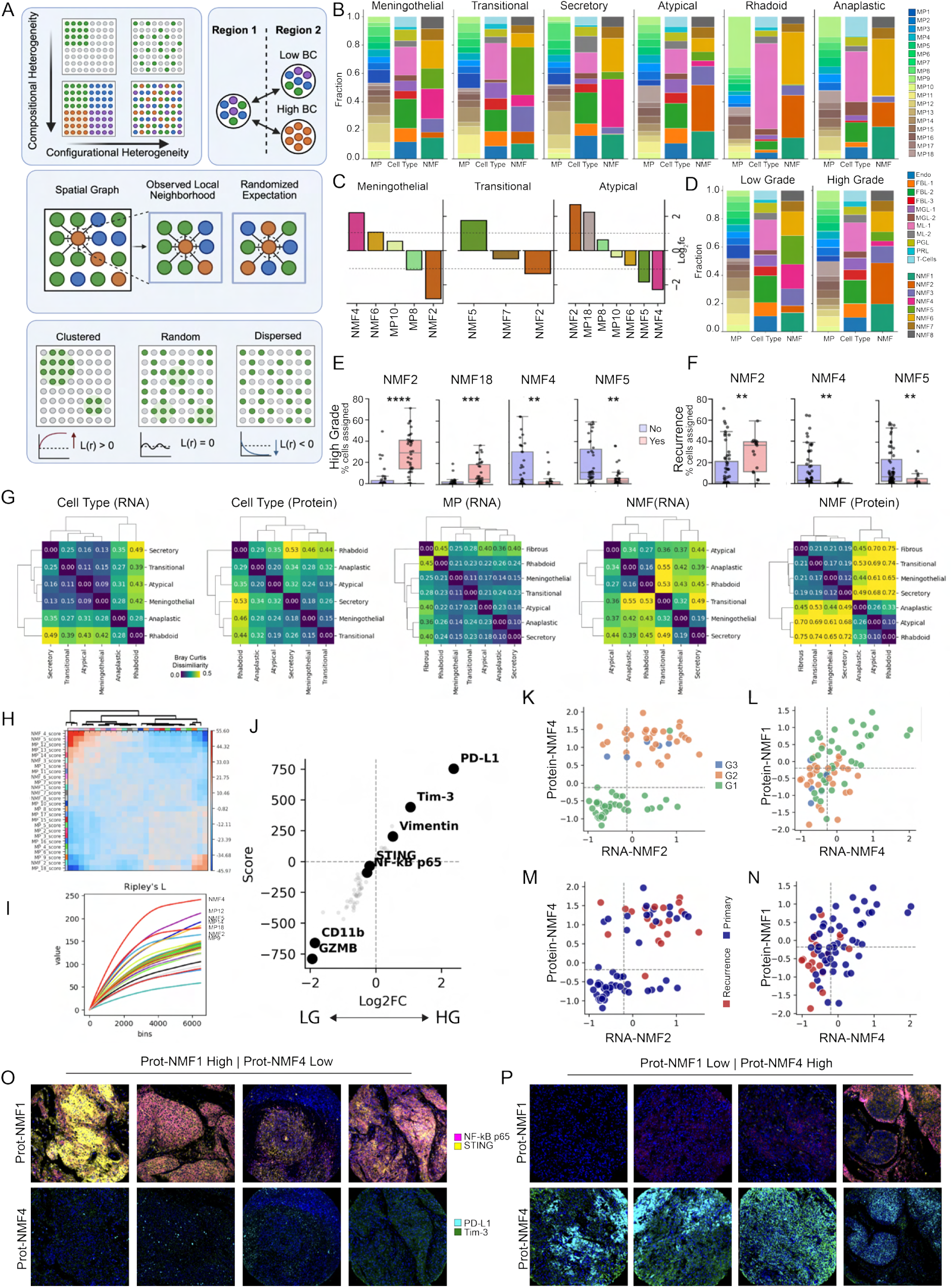
Spatial molecular programs uncover organized tumor states predictive of aggressive features. **A.** Schematic demonstrating spatial statistics terminology, including configurational and compositional heterogeneity, Bray-Curtis dissimilarity, Ripley’s L statistic, and neighborhood analysis. Created in BioRender. https://BioRender.com/ydlj6k6 **B.** Stacked bar plots showing the composition of cell types, RNA-NMF components, and RNA meta-programs (RNA-MPs), stratified by histology, including meningothelial (n = 25), transitional (n = 12), secretory (n = 4), atypical (n = 30), rhabdoid (n = 2), and anaplastic (n = 4). **C.** Log_2_ fold-change of significantly enriched components across histologic subtypes: meningothelial (n = 25), transitional (n = 12), and atypical (n = 30). **D.** Stacked bar plots showing the composition of cell types, RNA-NMF components, and RNA meta-programs (RNA-MPs), stratified by grade. **E.** Box-and-whisker plots showing the distribution of the dominant component proportion across individual patient samples, comparing low-grade and high-grade tumors (top), and **F.** Recurrence (vs. primary) status (bottom). Points denote aggregate tumor FOVs by individual; center lines indicate the mean, and whiskers represent ±1 standard deviation. * p-adj < 0.05, ** p-adj < 0.01, *** p-adj < 0.001, **** p-adj < 0.0001 **G.** Bray–Curtis dissimilarity indices summarizing intratumoral heterogeneity, grouped by (left to right) RNA-defined cell type, protein-defined cell type, RNA-defined meta-programs (RNA-MPs), RNA-defined NMF components (RNA-NMFs), and protein-defined NMF components (Protein-NMFs). **H.** Hierarchical clustering of spatial neighborhood co-occurrence analysis of RNA-NMF and RNA-MP components within spatial domains. **I.** Ripley’s L statistic showing spatial clustering (see A) for each component. **J.** Scatterplot showing log_2_ fold-change and score calculated by Scanpy differential expression analysis by grade (low-grade vs. high-grade); highlighted are the top genes for high-grade and low-grade tumors. **K–L.** Scatterplots showing RNA-NMF2 (oxidative phosphorylation) versus Protein-NMF4 (K, high-grade–associated) and RNA-NMF4 (vascular remodeling) versus Protein-NMF1 (L, low-grade–associated). Points are colored by WHO grade; dotted lines mark the 50th-percentile threshold. **M–N.** Scatterplots colored by recurrence status, showing RNA-NMF2 (oxidative phosphorylation)/Protein-NMF4 (M) and RNA-NMF4 (vascular remodeling)/Protein-NMF1 (N). **O-P.** Representative images showing localization of multi-modality data and spatial enrichment patterns for RNA-NMF2 (oxidative phosphorylation)/Protein-NMF4–enriched tumors **(O)** and RNA-NMF4 (vascular remodeling)/Protein-NMF1 (STING^+^/NF-*κ*B p65^+^)–enriched tumors **(P).**

Atypical tumors were characterized by enrichment of programs associated with aggressive tumor biology, including RNA-NMF2 (oxidative phosphorylation) (log_2_FC = 2.68, p_adj_ = 8.11 × 10^−7^; Figure 3 C), RNA-MP18 (hypoxia) (log_2_FC = 2.24, p_adj_ = 9.01 × 10^−3^), and MP8 (CXCL12/CXCR4 signaling pathway) (log_2_FC = 0.63, p_adj_ = 2.92 × 10^−2^). These tumors showed concomitant depletion of vascular- and stromal-associated programs, including RNA-NMF4 (vascular remodeling) (log_2_FC = –2.28, p_adj_ = 3.50 × 10^−2^) and RNA-NMF5 (EMT) (log_2_FC = –1.83, p_adj_ = 3.15 × 10^−3^). In contrast, meningothelial tumors were enriched for vascular programs, including RNA-NMF4 (vascular remodeling) (log_2_FC = 2.21, p_adj_ = 3.46×10^−3^) and RNA-NMF6 (vascular morphogenesis) (log_2_FC = 1.07, p_adj_ = 1.36 × 10^−2^), alongside decrease of RNA-NMF2 (oxidative phosphorylation) (log_2_FC = –2.81, p_adj_ = 1.54 × 10^−3^) and MP8 (CXCL12/CXCR4 signaling pathway) (log_2_FC = –1.12, p_adj_ = 9.01 × 10^−3^). Transitional tumors exhibited intermediate profiles, with relative enrichment of RNA-NMF5 (EMT) (log_2_FC = 1.76, p_adj_ = 1.54 × 10^−3^) and partial depletion of RNA-NMF2 (oxidative phosphorylation) (log_2_FC = –1.35, p_adj_ = 3.83 × 10^−2^).

On aggregate, high-grade tumors demonstrated significant overrepresentation of RNA-NMF2 (oxidative phosphorylation) (log_2_FC = 2.66, p_adj_ = 4.62 × 10^−8^), RNA-MP18 (hypoxia) (log_2_FC = 2.25, p_adj_ = 5.94 × 10^−4^), and RNA-MP9 (cell cycle/G2M) (log_2_FC = 0.93, p_adj_ = 0.022) (Figure 3 D, E, Supplementary File 8). Conversely, low-grade tumors were enriched for vascular, fibrotic, and ECM–associated programs, including RNA-NMF4 (vascular remodeling) (log_2_FC = −2.50, p_adj_ = 0.0015), RNA-NMF5 (EMT) (log_2_FC = −1.71, p_adj_ = 0.0014), RNA-MP12 (RNA regulation) (log_2_FC = −1.13, p_adj_ = 0.033), and RNA-MP1 (extracellular matrix remodeling) (log_2_FC = −1.28, p_adj_ = 0.036).

A similar pattern was observed when comparing recurrent versus primary tumors (Figure 3 F, Supplementary File 8). Recurrent tumors showed enrichment of RNA-NMF2 (oxidative phosphorylation) (log_2_FC = 1.08, p_adj_= 0.0066), alongside marked depletion of RNA-NMF4 (vascular remodeling) (log_2_FC = −3.93, p_adj_ = 0.0066) and RNA-NMF5 (EMT) (log_2_FC = −1.74, p_adj_ = 0.0066).

To assess whether SMP composition reflects histology beyond cell-type identity, we computed Bray–Curtis dissimilarity using SMPs, RNA-based cell types, and protein-based cell types. Hierarchical clustering of Bray–Curtis dissimilarities revealed that tumors of similar grade exhibited more consistent compositional profiles when represented by SMPs than when represented by RNA- or protein-based cell-type annotations, which failed to consistently group these morphologies (Figure 3 G, Supplementary File 8). This indicates that SMPs capture shared program-level composition across tumors within the same histologic and grade categories. However, compositional similarity alone does not determine the arrangement of these programs within tissue.

We therefore quantified spatial configuration directly using Ripley’s L-statistic and neighborhood co-occurrence analysis (Figure 3 H, I). Neighborhood co-occurrence revealed that SMPs form non-random, spatially coherent clusters that organize into two dominant configurations: (i) a vascular–fibrotic cluster comprising RNA-NMF4 (vascular remodeling), RNA-NMF5 (EMT), and RNA-MP12 (RNA regulation), and (ii) an aggressive cluster composed of RNA-NMF2 (oxidative phosphorylation), RNA-MP9 (cell cycle/G2M), and RNA-MP18 (hypoxia) (Figure 3 H). These components, together with RNA-MP14 (stress), exhibited the strongest spatial clustering by Ripley’s L-statistic (Figure 3 I).

PD-L1 and TIM-3 were among the most strongly overexpressed proteins in high-grade tumors and constituted defining markers of Protein-NMF4 (Figure 3 J, Supplementary File 8). Cross-modality analysis demonstrated strong correspondence between RNA- and protein-defined programs (Supplementary Figure S8 C). RNA-NMF2 (oxidative phosphorylation) showed a robust correlation with Protein-NMF4 (PD-L1^+^/TIM-3^+^; RNA-NMF2 (oxidative phosphorylation) r = 0.61, Figure 3 K, M). Co-enrichment of RNA-NMF2 (oxidative phosphorylation) and Protein-NMF4 (PD-L1^+^/TIM-3^+^) above the 50th percentile was strongly associated with high-grade and recurrence, exhibiting high sensitivity (0.83) and specificity (0.98) for high-grade tumors (Figure 3 K, Supplementary File 8), and sensitivity of 0.79 with specificity of 0.72 for recurrence (Figure 3 M).

In contrast, RNA-NMF4 (vascular remodeling) correlated with Protein-NMF1 (STING^+^; r = 0.53, Figure 3 L, N). Co-enrichment of RNA-NMF4 (vascular remodeling) and Protein-NMF1 (STING^+^) showed high specificity (0.89) for low-grade tumors (Figure 3 L) and for primary (non-recurrent) tumors (0.89) (Figure 3 N). The predominance of expression of either STING^+^ or PD-L1^+^/TIM-3^+^ within tissue sections was apparent at the protein level (Figure 3 O, P).

#### Identification of a molecular signature corresponding to whorls and lobules

Microanatomical structures known as whorls (concentric onion-skin-like arrangements of meningothelial cells) and lobules (rounded well-circumscribed nodules of tumor separated by fibrovascular septa) are hallmark features of meningioma histology, yet their molecular organization and relationship to dominant spatial tumor programs remain poorly defined. During initial exploratory analyses, unsupervised Leiden clustering was performed on individual Visium sections to identify spatially coherent transcriptional regions. While these clusters were not directly comparable across samples, several sections exhibited reproducible, compact spatial domains with shared gene expression features. Subsequent neuropathologic review identified these regions as containing meningioma whorls and lobules (Figure 4 A–C). Across these three whole-transcriptome–sequenced samples, this analysis identified a shared set of 28 genes consistently enriched within whorl- and lobule-associated regions (Supplementary File 7). Among the most highly expressed genes within the top 10 of each tumor were *NDUFA4L2*, *CCN3*, and *CCND1* (Figure 4 D–F). To enable quantitative analysis across the full cohort, we defined a continuous whorl–lobule (WL) score based on the median expression of these 28 genes. Gene set enrichment analysis of the WL score revealed pathways related to VEGF, Hippo, TGF*β* signaling, and glycolysis, suggesting coordination of metabolic, growth, and stress-response programs within these regions (Figure 4 G). Although the genes underlying the whorl–lobule score were derived from a limited number of Visium-profiled tumors, integration with SMPs enabled evaluation of WL–associated states across the full cohort.

**Figure 4.**
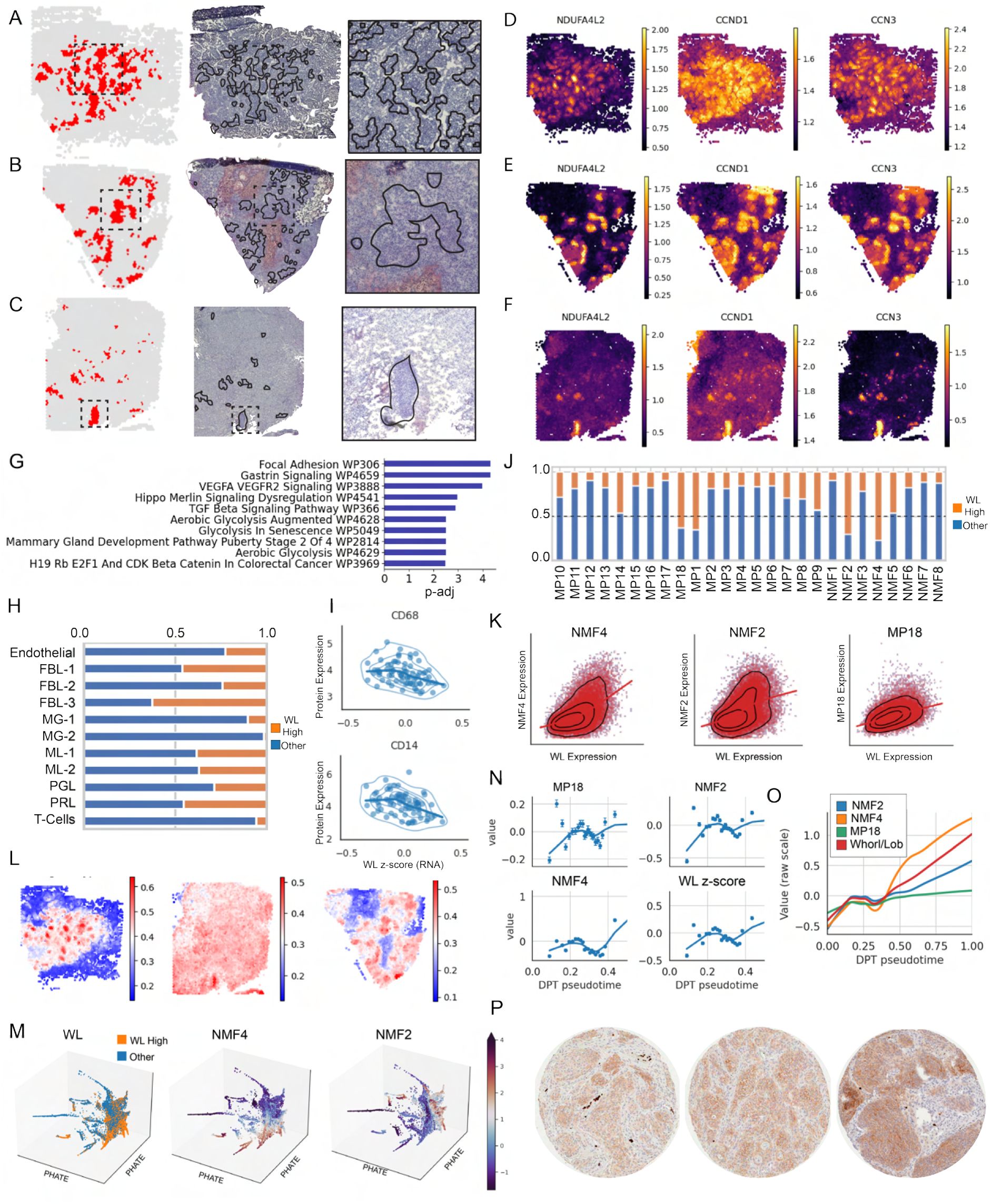
Whorl–lobule architecture links histology to hypoxia- and SMP-associated ecological states. **A–C.** Representative whole-section H&E images of meningioma tumors (Yale-NS-1206 (top), Yale-NS-0848 (middle), Yale-NS-0856 (bottom)), corresponding spatial maps of whorl/lobule (WL) signature enrichment inferred from spatial transcriptomics (left), spatially coherent whorl–lobule score–associated domains overlaid on histology (middle), and magnified regions (right). **D–F.** Spatial expression maps of top genes contributing to the whorl–lobule score across the tumors shown in **A–C**, demonstrating localized enrichment within whorl–lobule–associated regions. **G.** Gene set enrichment analysis of genes underlying the whorl–lobule score. **H.** Fraction of cells within each RNA-defined cell type scoring in the top 30th percentile of the whorl–lobule score. **I.** Patient-level correlations between the whorl–lobule score and protein abundance of CD14 and CD68. **J.** Fraction of cells assigned to each spatial molecular program (SMP) scoring in the top 30th percentile of the whorl–lobule score. **K.** Per-cell correlation density plots between the whorl–lobule score and key SMPs (RNA-NMF4 (vascular remodeling), RNA-NMF2 (oxidative phosphorylation), RNA-MP18 (hypoxia)), demonstrating continuous associations. **L.** Spatial expression of the MSigDB hypoxia gene set across the tissue sections shown in **A–C**, highlighting heterogeneity of hypoxic signaling within whorl–lobule–associated regions. **M.** PHATE embedding of single cells colored by the whorl–lobule score and expression of RNA-NMF4 (vascular remodeling), RNA-NMF2 (oxidative phosphorylation), and RNA-MP18 (hypoxia). **N.** Pseudotime analysis along the PHATE trajectory showing coordinated changes in the whorl–lobule score, RNA-NMF4 (vascular remodeling), RNA-NMF2 (oxidative phosphorylation), and RNA-MP18 (hypoxia) expression. **O.** Extended pseudotime trajectory illustrating divergence from RNA-NMF4 (vascular remodeling) toward RNA-NMF2 (oxidative phosphorylation) and RNA-MP18 (hypoxia) states, with cells exhibiting high whorl–lobule scores concentrated along this transition. **P.** HIF-1*α* immunohistochemistry showing nuclear staining within a subset of whorl–lobule structures.

Consistent with these pathway-level findings, the WL score was most enriched within FBL-1 (46.1%), FBL-3 (62.6%), and Proliferative cell populations (45.4%) (Figure 4 H, Supplementary File 7), cell types previously designated as tumor-associated and identified as active participants in WNT, BMP, and TGF*β* signaling (Supplementary Figure S4). In contrast, regions with high WL scores exhibited relative depletion of MG-1 (10.4%) and MG-2 (1.6%). This pattern was corroborated at the protein level by inverse correlations with CD14^+^ and CD68^+^ macrophages (Figure 4 I; Spearman *r* = –0.41, *p* = 2.69e-4 for CD14; *r* = –0.40, *p* = 3.14e-4 for CD68).

To contextualize the WL score within the broader spatial molecular landscape, we examined its overlap with SMPs. Cells in the top 30th percentile of WL score were disproportionately assigned to RNA-NMF4 (vascular remodeling) (76.9%), RNA-NMF2 (oxidative phosphorylation) (69.9%), RNA-MP1 (extracellular matrix remodeling) (64.4%), and RNA-MP18 (hypoxia) (62.6%) (Figure 4 J, Supplementary File 7). Given the previously established clinical relevance of RNA-NMF4 (vascular remodeling), RNA-NMF2 (oxidative phosphorylation), and RNA-MP18 (hypoxia), we further quantified their relationship to the WL score.

Across samples, WL scores correlated positively with RNA-NMF4 (vascular remodeling) (*r* = 0.52), RNA-NMF2 (oxidative phosphorylation) (*r* = 0.48), and RNA-MP18 (hypoxia) (*r* = 0.47), supporting a continuous association between these spatial programs and WL–associated regions (Figure 4 K). To explore the structure of this relationship, we embedded cells using PHATE followed by pseudotime inference (Figure 4 M-O). This analysis revealed a branching trajectory in which cells with high WL scores occupied an intermediate position between RNA-NMF4 (vascular remodeling)– and RNA-NMF2 (oxidative phosphorylation)–dominated states, with the RNA-NMF2 (oxidative phosphorylation) branch following an RNA-MP18 (hypoxia) trajectory, consistent with earlier analyses linking EMT and hypoxia-associated programs (Figure 4 L, N–O).

To assess hypoxia directly, we performed immunohistochemical staining for HIF-1*α* (Figure 4 P). HIF-1*α* expression localized specifically to a subset of regions with high WL scores, but was not uniformly present across all WL structures, indicating the existence of molecularly distinct WL subtypes, including both hypoxic and non-hypoxic variants.

Because spatial profiling is costly and limited in scale, we next asked whether the SMPs discovered here can be decoded directly from routine H&E, thereby extending a discovery atlas into a scalable measurement framework. While spatial molecular programs capture these organized tumor states and associate with aggressive features, their discovery is inherently limited to tissues profiled by spatial transcriptomics. This constraint restricts both clinical translation and the ability to systematically relate expression-defined programs to microanatomical structures across large cohorts. We therefore sought to determine whether these spatial molecular programs and their associated WL states, as quantified by the WL score, could be inferred directly from routine histopathology, thereby enabling extension to whole-slide images (WSIs) and independent datasets.

### A generalizable and interpretable hybrid deep learning model predicts spatial molecular programs and clinical stratification

Spatial profiling remains costly, technically demanding, and unavailable in routine clinical practice, whereas H&E histopathology is universally available. To determine whether SMP architecture could be recovered from routine pathology specimens alone, we developed MeningNet, a hybrid ConvNeXt–Vision Transformer with component-specific attention pooling, to predict spatial molecular program scores directly from H&E patches while preserving spatial interpretability at the tissue level (Figure 5). This approach enables molecular inference in annotation-free sections, extends the SMP inference to large external cohorts lacking matched spatial transcriptomic data, and, in principle, could support deployment in cohorts lacking molecular profiling, such as in resource-limited settings where molecular profiling is not routinely available.

**Figure 5.**
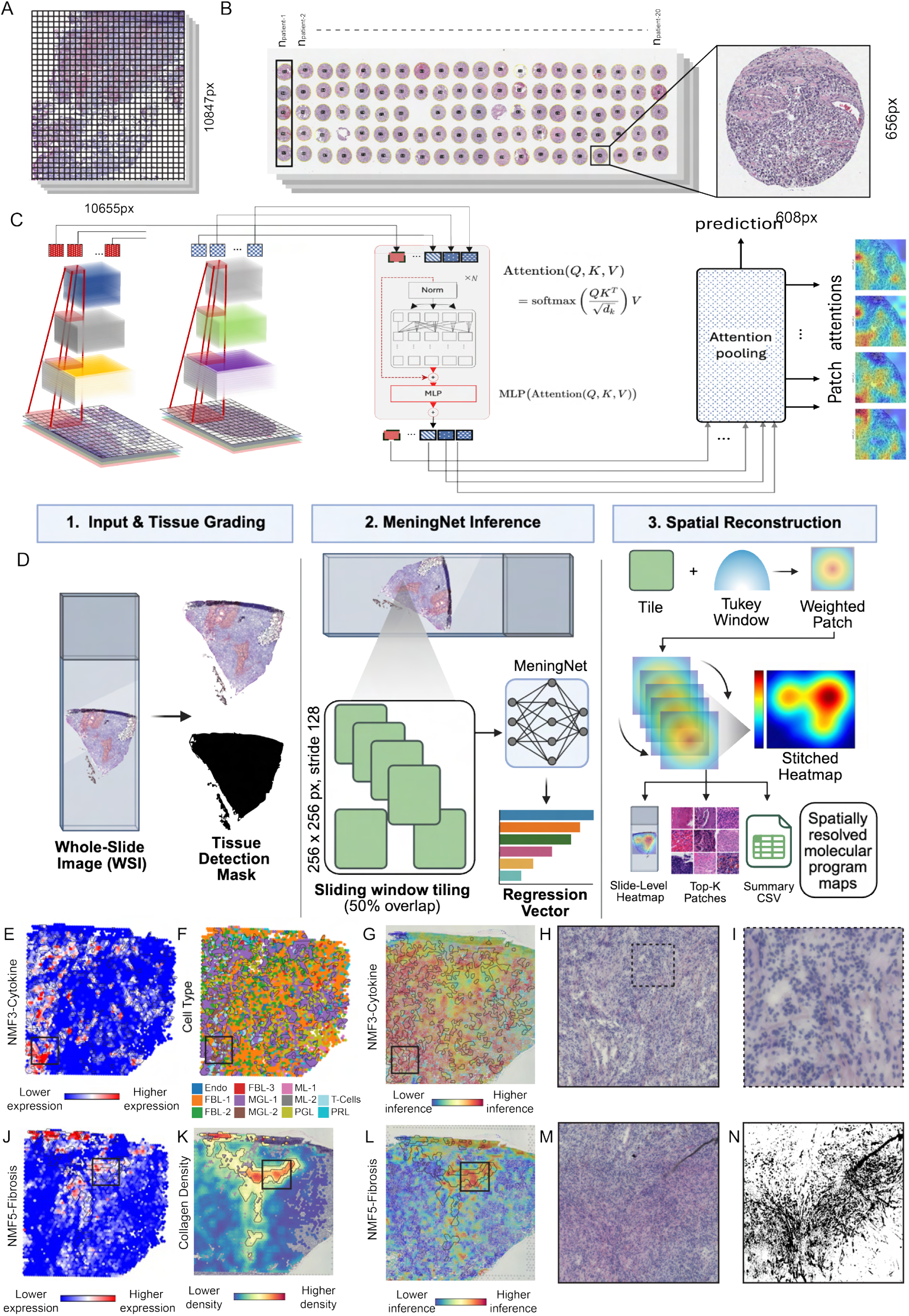
MeningNet decodes spatial molecular programs from routine H&E. **A-B.** Schematic overview of general layouts of H&E belonging to Visium (A) and NanoString CosMx (B) spatial transcriptomic data **C.** Architecture of MeningNet, a hybrid ConvNeXt–Vision Transformer model that predicts spatial molecular program scores directly from H&E image patches. Convolutional feature extraction captures local cytoarchitectural patterns, while transformer-based attention pooling models broader spatial context. The model outputs continuous program scores and generates component-specific attention maps to enable biological interpretation. **D.** Schematic of the sliding-window MeningNet workflow for whole-slide inference and spatial reconstruction. Whole-slide images are segmented to identify tissue regions, tiled using a sliding-window strategy, and processed by MeningNet to generate per-patch molecular predictions that are aggregated into stitched, spatially resolved score heatmaps. Created in BioRender. https://BioRender.com/7a0vq9o **E-N.** Representative examples demonstrating correspondence between biological ground truth and sliding-window MeningNet inference in Visium-profiled meningiomas (Yale-NS-0691). Predicted spatial maps recapitulate immune-associated (RNA-NMF3) and EMT-associated (RNA-NMF5) programs and localize to histologically consistent regions across scales. **E-I.** Immune-associated component. **E.** Whole-section heatmap showing spatial transcriptomic expression of RNA-NMF3 (cytokine/immune); boxed region denotes area of interest. **F.** Cell-type integration using CytoSPACE, highlighting immune cell localization. **G.** Sliding-window MeningNet inference and stitched score heatmap for RNA-NMF3. **H.** Magnified view of H&E section of region of interest. **I.** Further magnification highlighting local cellular morphology. **J-N.** RNA-NMF5 (EMT). **J.** Whole-section heatmap showing spatial transcriptomic expression of RNA-NMF5 (EMT); boxed region represents ROI. **K.** Collagen density map computed from the corresponding H&E image. **L.** Sliding-window MeningNet inference and stitched score heatmap for RNA-NMF5 (EMT). **M.** Magnified H&E image of the boxed region. **N.** Magnified H&E binary mask highlighting collagen-rich architecture.

MeningNet was trained to regress SMP scores directly from H&E image patches while generating component-specific attention maps to provide spatial interpretability at the tissue level. To assess generalizability, we applied sliding-window inference and spatial reconstruction across entire tissue sections, followed by patient-level aggregation of patch-level predictions to evaluate CNS WHO grade discrimination (Figure 5 D). Performance was validated in an internal replication cohort (n = 648 fields of view (FOVs), 128 tumors) and in an external cohort from the Digital Brain Tumor Atlas (n = 465 WSIs, Figure 6 A).^38^ Across cohorts, MeningNet recapitulated spatial molecular programs and enabled grade stratification without retraining.

**Figure 6.**
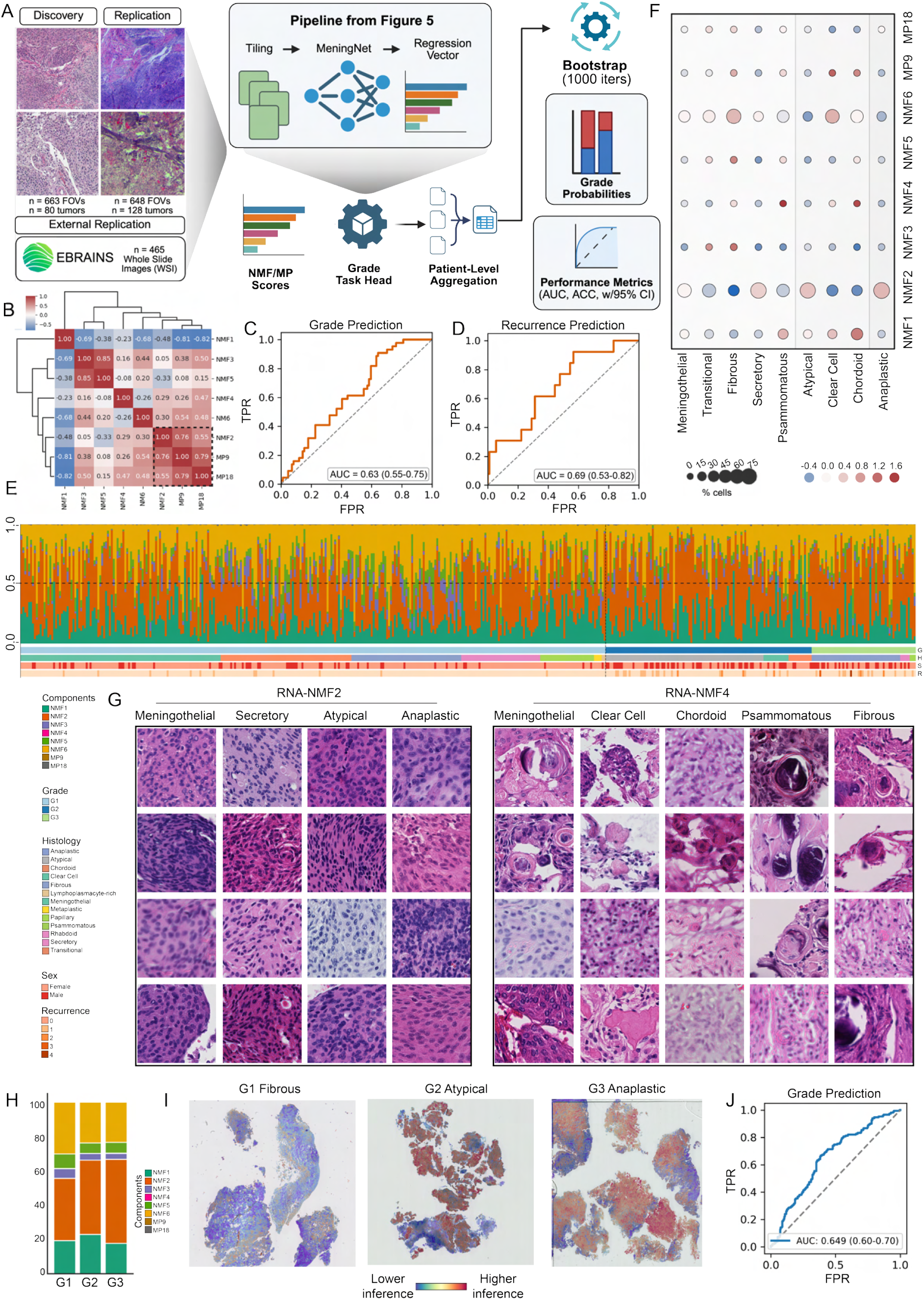
MeningNet generalizes across internal replication and external DBTA cohorts. **A.** External validation workflow. DBTA whole-slide H&E images were tiled and processed using the pretrained MeningNet pipeline (Figure 5) to generate patch-level spatial molecular component predictions, which were aggregated at the patient level for grade and recurrence inference with bootstrap-based performance estimation. Created in BioRender. https://BioRender.com/1ex74bd **B.** Hierarchically clustered correlation matrix of inferred component scores across DBTA tumors, recapitulating separation of aggressive (RNA-NMF2 (oxidative phosphorylation), MP9 (cell cycle/G2M), MP18 (hypoxia)) and immune/fibrotic (RNA-NMF3 (cytokine/immune), RNA-NMF5 (EMT)) components. **C.** ROC curves for discrimination of low-grade (G1) versus high-grade (G2/G3) and **D.** 5-year post-op recurrence prediction. **E.** Patch-level dominant component distributions across n = 465 DBTA samples. **F.** Cohort-level pseudo-spatial enrichment of RNA-NMF components across DBTA samples. **G.** Representative high-scoring patches for RNA-NMF2 (oxidative phosphorylation) and RNA-NMF4 (vascular remodeling), illustrating aggressive hypercellular versus laminated/calcified architecture. **H.** Distribution of dominant pseudo-spatial components per tumor, stratified by grade. **I.** Representative whole-slide examples of RNA-NMF2 (oxidative phosphorylation) in tumors across WHO grade. **J.** Overall grade discrimination performance with bootstrapped 95% confidence interval (n = 465).

#### MeningNet predicts spatial molecular programs from histology in expression-matched sections

MeningNet, a hybrid ConvNeXt–Vision Transformer architecture with attention pooling, was trained on a mixed-modality dataset comprising Visium-profiled fresh-frozen sections and adjacent H&E sections corresponding to NanoString CosMx FFPE samples (Figure 5 A–C).

Using a multi-task framework, a shared image backbone extracted multiscale features that were passed to task-specific MLP heads to regress eight spatial molecular programs (SMPs): six clinically enriched components (RNA-NMF1, RNA-NMF2 (oxidative phosphorylation), RNA-NMF4 (vascular remodeling), RNA-NMF5 (EMT), MP9 (cell cycle/G2M), MP18 (hypoxia)) and two components with established morphologic correlates (immune-associated RNA-NMF3 and RNA-NMF6 (vascular morphogenesis)). Unlike a single global slide embedding, MeningNet learns one attention-pooled representation per SMP, enabling distinct spatial evidence patterns to support different molecular programs within the same tissue section. Auxiliary grade and recurrence heads were used during representation learning, whereas the clinical performance reported here was evaluated from slide-level aggregation of predicted SMPs unless otherwise stated.

For Visium samples, 56,020 spatial spots were paired with matched 256 × 256 pixel H&E patches. For CosMx samples, FOV SMP scores were computed by aggregating cell-level expression and subdividing each image into 4 patches, yielding 2,588 image–expression pairs.

Using patient-stratified splits, the model achieved robust held-out regression performance (mean Spearman r = 0.67, mean Pearson r = 0.72, mean R^2^ = 0.48). Per-component Spearman correlations ranged from 0.29 (MP9 (cell cycle/G2M)) to 0.78 (RNA-NMF4 (vascular remodeling)) (Supplementary Figure S10 A).

In Visium-matched sections, inferred SMP maps recapitulated expected spatial patterning. RNA-NMF3 (cytokine/immune) localized to cytokine-rich immune niches and aligned with CytoSPACE-derived immune cell localization (Figure 5 E–I). RNA-NMF5 (EMT) localized to collagen-dense perivascular regions, confirmed by independent collagen density maps and high-magnification histology (Figure 5 J–N). Sliding-window inference also reproduced RNA-NMF2 (oxidative phosphorylation) enrichment in WL-associated regions defined by gene expression (Supplementary Figure S10 C).

#### Spatial molecular programs generalize across preparations

We next applied MeningNet to an internal replication cohort comprising independently prepared H&E sections (n = 648 FOVs, 128 tumors) imaged using distinct microscopy platforms, and to an external replication cohort consisting of 520 publicly available WSIs from the Digital Brain Tumor Atlas (DBTA), of which 465 remained after post hoc exclusion of underrepresented histologic subtypes (Figure 6A).^38^ In the internal replication cohort, hierarchical clustering of image-derived SMP correlation scores recapitulated relationships observed in the transcriptomic data, including co-occurrence of RNA-NMF2, enriched for oxidative phosphorylation, with MP9, enriched for cell cycle/G2M programs, and MP18, enriched for hypoxia-associated programs (Figure 6B). At the patient level, aggregation of patch-level SMP predictions showed significant but modest association with WHO grade (AUC = 0.63, 95% CI: 0.55–0.75; 1,000 bootstrap resamples) and five-year recurrence (AUC = 0.69, 95% CI: 0.53–0.82; top 10% patch aggregation; 1,000 bootstrap resamples; Figure 6C-D). These findings are consistent with the framework capturing risk-relevant biology beyond what is reflected in WHO grade alone, although with limited stand-alone predictive performance.

Separately, the DBTA evaluation assesses whether morphology–SMP relationships learned in paired sections remain stable under external domain shifts, including changes in cohort composition and imaging conditions. Without retraining, MeningNet was applied to our external cohort from the DBTA, encompassing the full spectrum of WHO-recognized meningioma histologies (Figure 6 E, Supplementary Figure S11 A, Supplementary File 9). Sliding-window inference generated spatially resolved SMP maps across all slides. Aggregation of patch-level predictions revealed differential component usage across histologic subtypes, including enrichment of RNA-NMF2 (oxidative phosphorylation) in atypical and anaplastic meningiomas and enrichment of RNA-NMF4 (vascular remodeling) in psammomatous, chordoid, fibrous, and clear cell tumors (Figure 6 F, Supplementary File 9).

RNA-NMF4 (vascular remodeling) signal was observed within subsets of WL structures and localized prominently to psammoma bodies across stages of maturation (Figure 6 G). Notably, this spatial distribution parallels the transcriptomic continuum identified by PHATE analysis (Figure 4 N–O), in which RNA-NMF4 (vascular remodeling) occupies a trajectory branch diverging from RNA-NMF2 (oxidative phosphorylation). These findings demonstrate concordance between RNA-defined program structure and morphology-based inference across both non-calcified WL-associated regions and progressively calcified psammomatous domains. When comparing between CNS WHO grade, we observed that RNA-NMF2 (oxidative phosphorylation) was identified in a higher proportion in G2 and G3 meningiomas (Figure 6 H, I).

Using patient-level aggregation, grade prediction across all DBTA samples yielded an AUC of 0.61 (95% CI: 0.56–0.67, mean with bootstrapping = 1000), sensitivity 0.68, and specificity 0.54. Error analysis revealed high (> 0.60) error rates in angiomatous and microcystic meningiomas, histologic subtypes not represented in the training dataset (Supplementary Figure S11C, Supplementary File 9). Excluding these underrepresented subtypes (n = 55) improved performance to an AUC of 0.65 (95% CI: 0.60–0.70, mean with bootstrapping = 1000), sensitivity 0.67 (95% CI: 0.60–0.83), and specificity 0.62 (95% CI: 0.43–0.69) (Figure 6 J).

## Discussion

Our data support a model in which meningioma-level molecular continua arise from the differential representation and spatial organization of discrete, reproducible tissue states embedded within canonical microanatomy, rather than from unstructured gradients. In this study, we show that meningiomas are organized as modular ecosystems composed of such states, which we term spatial molecular programs (SMPs). These findings position tumor architecture as an organized, modular system encoding clinically relevant biology.

This study defines SMPs as reproducible, multicellular architectural states that organize meningioma tissue across patients, histologic subtypes, and experimental platforms. SMPs capture coordinated transcriptional configurations spanning malignant, immune, and stromal compartments that recur as spatially coherent tissue units, integrating and extending isolated cell-type signatures. These units are conserved across fresh-frozen and FFPE specimens and across independent spatial profiling technologies, and are recapitulated by deep-learning inference from routine histology in held-out and external cohorts. Importantly, SMPs are not restricted to specific histologic categories but are differentially utilized across tumors, suggesting that morphologic diversity reflects variable composition of a limited set of stable architectural building blocks.

Recent integrative analyses in meningioma have emphasized tumor-level molecular continua shaped in part by microenvironmental composition.^22^ Our findings provide architectural resolution to this concept by demonstrating that such continua arise from reproducible tissue states, rather than unstructured gradients. In this compositional model, tumors deploy shared histopathological configurations in varying proportions and arrangements, reconciling continuum-like transcriptomic patterns with stable microanatomical structures.

A central organizing principle emerging from this atlas is the presence of spatial convergence zones in which metabolic stress, extracellular matrix remodeling, vascular signaling, and proliferative programs co-localize in reproducible configurations. These states recur across patients and grades, with subsets encoding components strongly associated with clinical aggressiveness, suggesting that meningioma tissue operates within modular architectures.

A defining histologic feature of meningioma is the presence of meningothelial whorls and lobules.^17^ Here, we demonstrate that whorl- and lobule-associated regions correspond to transcriptionally structured spatial compartments independently identified by spatial profiling and deep-learning inference from routine histology. Rather than representing uniform entities, these architectural units span a structured continuum of molecular states.

Along one branch of this continuum, RNA-NMF2 is enriched for hypoxia-, metabolic stress-, and EMT–associated programs and is characterized by elevated PD-L1 expression with relative immune exclusion. This configuration is preferentially observed in tumors with aggressive histopathological features, suggesting that a subset of whorl-associated regions occupies a metabolically constrained, immune-evasive phenotype. Prior studies have linked PD-L1 expression to higher-grade meningioma,^39^ and ultrastructural work has shown that whorl formation can be induced under nutrient deprivation and is associated with enhanced membrane polarization and intercellular junctions,^40^ consistent with coordinated metabolic adaptation.

In contrast, RNA-NMF4 defines a distinct axis characterized by extracellular matrix remodeling, vascular signaling, ion-handling pathways, and STING/NF*κ*B activation, and spatially corresponds to progressively calcified psammomatous regions associated with more indolent behavior. Prior ultrastructural studies describe a stepwise mineralization process culminating in laminated psammoma bodies.^41^ The enrichment of STING/NF*κ*B signaling within this axis suggests that psammomatous architecture arises within a chronically organized, matrix-remodeling niche.

Low-dimensional embeddings and trajectory analyses recover these states as branching structures rather than isolated clusters, and deep-learning inference from routine histology independently reconstructs these architectural gradients. Together, these findings support a model in which canonical meningioma architecture defines a continuous yet structured ecological landscape containing discrete molecular basins, analogous to attractor states within a Waddington-like framework.

The value of MeningNet lies not in moderate prediction of grade or recurrence alone, but in its ability to produce spatially resolved, biologically interpretable latent variables from routine histology, extending spatial multi-omic discovery into a scalable assay. Inferred spatial features enabled discrimination of tumor grade and recurrence risk in both internal and external cohorts, and tumors discordant with grade classification frequently exhibited elevated RNA-NMF2 abundance. This suggests that morphology-derived SMPs capture biologically meaningful risk information not fully represented by conventional CNS WHO grading.

Importantly, this study isolates the contribution of spatial architecture without incorporating genomic or epigenomic features. DNA methylation classifiers and recurrent driver mutations provide complementary tumor-intrinsic information, whereas SMPs capture the ecological context in which these alterations are expressed. Integration of SMPs with genomic and epigenomic profiling may enable architecture-aware classification systems that jointly model tumor-intrinsic alterations and microenvironmental organization.

Several limitations warrant consideration, including the under-representation of CNS WHO grade 3 tumors and the cross-sectional nature of the analysis. While compositional and spatial analyses indicate that SMPs are not reducible to cell-type composition, formal statistical tests of this emergence—and demonstration that this principle generalizes across tumor types—remain important directions for future work. Because external grade discrimination remained modest and performance degraded in histologic subtypes not present in the training set, MeningNet should be interpreted as a biological decoder of ecological state rather than a stand-alone diagnostic classifier. Additionally, some SMPs (e.g., MP9 and MP18) exhibited weaker performance, likely reflecting limited training data. Future longitudinal and perturbation studies will be required to define the mechanisms governing spatial state formation and transitions.

Together, these findings support a model in which meningiomas are organized as modular ecosystems whose architecture encodes molecular and clinical information accessible from standard pathology.. The ability to decode spatial molecular states directly from routine histology provides a scalable bridge between spatial biology and clinical practice and establishes a foundation for multimodal, architecture-aware precision diagnostics.

## METHODS

### Sample Collection and Preparation

Tumor specimens were obtained with written informed consent from all participants under Institutional Review Board–approved protocols (HIC # 9406007680) at Yale New Haven Hospital. All samples were reclassified according to the WHO CNS 2021 guidelines by a neuropathologist (D.M.).^17^ In total, 141 patients across meningioma grades 1-3, histologic subtypes, and genomic backgrounds were included (Supplementary File 1). Among the 23 scRNA samples, 10 samples from 6 patients were obtained from publicly available data.^19^ Clinical data (patient demographics, tumor location, histopathological diagnosis, and follow-up outcomes) were recorded for each sample (Supplementary File 1). Additionally, representative normal meningeal/dural tissue from 3 patients was obtained as a non-tumor reference. Samples included in the TMA previously underwent bulk DNA Whole-exome sequencing to characterize genomic alterations.^15^ Determination of mutation subgroup was analyzed using previously published methods from the Günel lab.^11^ Sex was recorded for all donors (Supplementary File 2) and reflects the established female-predominant sex distribution of meningioma. The influence of sex on the composition of SMPs was not formally evaluated in this study.

#### Tissue Preparation for Spatial Samples

**Visium 10x** Fresh tumor tissues were collected in the operating room and either processed immediately for single-cell sequencing or flash-frozen in OCT and stored at −80°C for spatial transcriptomic profiling. To prepare for use with 10x Visium Spatial Sequencing, OCT blocks were sectioned into 10µm slices and immediately placed on Visium array slides (Visium Spatial Gene Expression slides, 10× Genomics). Hematoxylin and Eosin (H&E) staining, RNA extraction, and cDNA strand synthesis were followed as directed in the Visium protocol. Images were captured using the Keyence BZ-X700 microscope system using 40x magnification. Sequencing was performed at Yale Center for Genome Analysis, and Visium samples were sequenced to 75 million pairs/sample. Transcriptome alignment and counts were generated using 10x Space Ranger version 2.1.0.

**CosMx NanoString** A tissue microarray was constructed from 119 genetically and clinically annotated meningiomas preserved as formalin-fixed paraffin-embedded (FFPE) specimens. For each tumor, five representative regions of interest were selected by D.F.M. in conjunction with a neuropathologist (D.M.). Samples were processed at the Yale Center for Genome Analysis using the NanoString CosMx Spatial Molecular Imager (SMI) platform for human 6K Discovery panel for multiplexed spatial RNA profiling (n = 77 tumors) and human immuno-oncology panel for spatial protein profiling (n = 119). Image processing was performed using the NanoString AtoMx Spatial Informatics Platform (CosMx Data Analysis v2.2; study version 1.3). Single-cell segmentation was performed using the default CosMx “Initial Segmentation” pipeline. Quality control was applied using per-cell fluorescence intensity metrics, including mean fluorescence intensity (MFI), intensity distribution percentiles (10th/90th), and IgG control signal to exclude low-quality cells. Protein expression values were normalized using the CosMx normalization module with per-cell intensity scaling and background correction based on control probes, followed by model-based estimation of per-cell protein abundance from raw fluorescence signals.

#### Tissue Preparation for Single-Cell RNA-seq

For both frozen and fresh tissue, samples were subjected to mechanical and enzymatic breakdown using the Brain Tumor Dissociation Kit by Miltenyi Biotec (#130-095-942). The solution consisted of 50 *µ*L of Enzyme N, 3890 *µ*L of Buffer X, 40 *µ*L of Buffer Y, and 20 *µ*L of Enzyme A. It was warmed to 37°C during sample preparation. The samples were then transferred to gentleMACS™ M Tubes (130-093-236) with Brain tumor lysis solution. They were processed using the gentleMACS dissociator program h_tumor_01, followed by rotation at 37°C for 10 minutes, and then processed with h_tumor_02 and rotated for 15 minutes at 37°C. Samples were washed with a 0.04% BSA/PBS mixture 1x before proceeding to either single-cell or single-nucleus-specific protocols.

**Fresh samples** After mechanical disruption, a 1x RBC lysis solution (RBC Cell Lysis Solution, Miltenyi Biotec 130-094-183) was prepared, and samples were incubated for 5 minutes at room temperature. The cells were washed with a 0.04%BSA/PBS mixture 2-3 times and filtered through a 45*µ*m filter. The samples were prepared with a target of 5,000-10,000 cells per sample, at a concentration of approximately 1200 cells/*µ*L in 50 *µ*L of 0.04% BSA/PBS, and sent to YCGA for library preparation.

**Frozen samples** Nuclei isolation was applied to frozen specimens. If specimens were in a block of OCT, the OCT was trimmed, and the sample was placed into a 1.5mL Eppendorf tube. A pestle was used to lyse the tumor while on dry ice. Once lysed into small pieces (grain size), the samples were washed with a 0.04 %BSA/PBS mixture 3 times and spun down at 300 rcf for 5 minutes. The cells were then lysed on ice for 5 minutes using the lysis buffer detailed below. Nuclei were washed once with the nuclei wash buffer and filtered through a 45 *µ*m filter. The nuclei were suspended in 10 million cells/1 mL and 50*µ*L of 7-AAD (eBioscience™ 00-6993-50), then incubated for 15 minutes at RT before FACS sorting for 7-AAD+ nuclei. The sorted nuclei were placed into 0.04%BSA/PBS solution, and a 10*µ*L aliquot was stained with fluorescent DAPI and inspected for quality before proceeding to library preparation by YCGA. *Cell Lysis Buffer Preparation* The lysis buffer was composed of various components, including: Tris-HCl (pH 7.4) at 10 mM, NaCl at 10 mM, MgCl2 at 3 mM, Bovine Serum Albumin (BSA) at 1%, Tween-20 at 0.1%, Dithiothreitol (DTT) at 1 mM, RNase Inhibitor at 1 U/*µ*L, Nonidet P40 Substitute at 0.1%, Digitonin at 0.01%, and Nuclease-Free Water at 1.67 mL. This combination facilitated cell lysis and preserved biological samples for further analysis. *Nuclei Wash Buffer Preparation* The nuclei wash buffer was prepared with the following components: Tris-HCl (pH 7.4) at 10 mM, NaCl at 10 mM, MgCl2 at 3 mM, BSA at 1%, Tween-20 at 0.1%, DTT at 1 mM, RNase Inhibitor at 1 U/*µ*L, and Nuclease-Free Water at a total volume of 3.40 mL.

#### Library Preparation and Sequencing for Single-Cell RNA-seq

Meningiomas were derived from three sources: Magill *et al.*,^19^ our frozen tissue repository, and prospectively collected fresh samples. For the samples we generated, sequencing was performed on a NovaSeq6000 system with paired-end sequencing at a depth of 250 million reads. The aligned transcriptome was processed using the GRCh38-2020-A reference. Our target range for cells was 5-10,000 cells per sample (Supplementary File 1).

### Data Processing and Quality Control

#### Single-cell RNA-seq

Raw single-cell RNA sequencing (scRNA-seq) data were processed using the 10x Genomics Cell Ranger pipeline (cellranger-7.1.0, GRCh38-2020-A). Downstream preprocessing was performed using Scanpy (v1.9). Cells were filtered based on library complexity and mitochondrial content, excluding cells below the 10th percentile and above the 90th percentile of total counts or with > 40% mitochondrial gene expression. Genes detected in fewer than 5 cells were excluded. Filtered datasets from individual samples were combined into a unified AnnData object, which was converted to Seurat. Batch correction was performed in Seurat (v5.1.0) using Harmony (v1.2.1) implemented in R (v4.4.1). After conversion of filtered AnnData objects to Seurat format, per-cell metadata, including sample identity and processing batch, were incorporated. Data were normalized, log1p transformed, and scaled. Principal component analysis (PCA) was performed on the variable-feature matrix using default parameters. Harmony was applied to the PCA embedding with the sequencing batch specified as the integration variable. The Harmony-corrected embedding was used for downstream analyses.

#### RNA Spatial transcriptomics

Spatial transcriptomic data generated using 10x Visium and NanoString CosMx platforms were processed as a combined dataset for joint analyses and as individual datasets for visualization and scRNA-seq integration. For Visium data, gene expression matrices aligned to histological images were generated using Space Ranger (v2.1.0). Spots were filtered per sample to exclude low-quality observations based on total counts (below the 5th percentile) and mitochondrial content (> 40%), and genes detected in fewer than 5 spots (Visium) or cells (CosMx) were removed. For CosMx RNA data, cell-level expression matrices and spatial metadata were imported into AnnData format. Quality control was performed per sample by visual inspection to filter out cells below the 10th percentile of total counts and detected genes, and to retain genes expressed in at least 5% of cells. After library normalization, control probes (e.g., “Negative” and “SystemControl”) were excluded from downstream analyses. For both platforms, expression matrices were log 1p transformed. To improve signal-to-noise ratio while preserving spatial structure, expression values were further processed using a graph-based diffusion framework (spARC),^42^ which integrates transcriptional similarity and spatial proximity to denoise expression values. Denoised and non-denoised post-filtered matrices were retained for downstream analyses.

For combined analyses, due to the difference in transcripts sequenced between datasets generated using 10x Visium and NanoString CosMx platforms, post-QC-filtered expression matrices with shared gene space were independently library-normalized, log 1p-transformed, and scaled. The normalized datasets were concatenated into a unified expression matrix and batch-corrected using HarmonyPy (v0.0.10) applied to the PCA embedding, with dataset batch (e.g., Visium batch or NanoString batch) and sequencing batch specified as the integration variable.

#### Protein Spatial transcriptomics

Single-cell spatial proteomics data were generated using the NanoString CosMx™ Spatial Molecular Imager (SMI) platform. The resulting protein expression matrices (stored as .h5ad files) were processed using Scanpy (v1.10.4). Initial quality control (QC) was performed at the sample level, assessing total protein signal per cell (nCount_RNA), the number of unique proteins detected per cell (nFeature_RNA), and total counts per field of view (FOV). To ensure robust downstream analysis, cells falling below the 5th percentile for total counts were excluded to mitigate noise from low-signal observations. Features were filtered to retain only those expressed in at least 1% of the total cell population. Following filtering, expression values were normalized using the Scanpy default normalize_total (scaling each cell to the median total counts of the dataset) and log-1p transformed.

#### Cell type annotation and spatial integration

We performed unsupervised Leiden clustering using Scanpy t-test_overestim_var with a resolution of 0.5 on post-processed, scRNA-seq data. We used Pearson correlation to discern differences between these clusters, setting a threshold of Pearson *r* > 0.5 for cluster amalgamation. This process yielded 11 different cell types that were annotated based on the enrichment of terms found in the scRNA-seq databases PanglaoDB, Tabula Sapiens, and CellMarkerAugmented, complemented by a manual review of the published meningioma scRNA-seq literature.^20,21^ Next, each cell was assigned a cell type label based on cluster identification, and Ecotyper was used to generate a high-confidence differential expression list and final assignment of cells to a cell type annotation matrix. To map single-cell–defined cell types onto spatial transcriptomic data, we applied CytoSPACE (v1.0.6) and spatial expression matrices derived from 10x Visium and NanoString CosMx. Mapping was performed under a honeycomb geometry using -sss consistent with Visium spot architecture, with an average of five cells assigned per spot and using -noss for NanoString CosMx, designed for single-cell spatial data.

Copy number variation (CNV) inference was performed using InferCNV (v1.28.0).^43^ Post-processed files were used as input, and genes were ordered according to genomic position based on the GRCh38 annotation. Cells were binned by previously established cell-type annotations. Analysis was performed with reference to “normal” dura. InferCNV was run using default parameters (smoothing window length of 101, cutoff value of 0.1) to denoise expression signals. CNV profiles were generated by averaging expression across adjacent genomic regions, with regions centered relative to the reference population.

### Defining spatial molecular programs

**Non-Negative Matrix Factorization.** Non-negative matrix factorization (NMF) was applied to the batch-corrected, normalized, and log-transformed expression matrix to identify continuous transcriptional programs shared across datasets. Prior to factorization, the expression matrix was linearly shifted non-negative values. Model fitting was performed across a range of factorization ranks (k = 2–11 for protein data and k = 2–21 for RNA data) using a subsampled set of up to 10,000 observations, NNDSVD initialization, and a maximum of 500 iterations. The optimal rank was selected based on the first local minimum in inter-component Jaccard overlap of top-ranked genes (50 genes for RNA; 5 for protein), corresponding to the onset of increasing redundancy (Supplementary Figure S9 A, B). Using the selected rank, NMF was refit on the full dataset. The resulting components represent latent transcriptional programs, with each observation assigned a vector of component scores. Scores were standardized across observations using z-score normalization, and the dominant program for each observation was defined as the component with the highest standardized score. Gene loadings were extracted and scaled to facilitate the interpretation of program-specific signatures.

#### Metaprogram derivation and projection

Gene programs were derived independently per sample using Leiden clustering (resolution = 0.5) and non-negative matrix factorization (NMF) across multiple factorization ranks (k = 2–10) and represented as gene sets, following previously described consensus program frameworks.^44^ Programs were filtered to retain signals that were reproducible across samples. Program similarity was quantified using Jaccard overlap, and programs sharing substantial gene overlap (Jaccard index > 0.2) were organized into a similarity graph, with nodes representing programs and edges indicating shared genes. Metaprograms (MPs) were defined using graph-based community detection to capture recurrent gene programs conserved across samples. For each MP, a consensus gene signature was constructed by ranking genes according to their frequency across constituent programs, and the top 50 genes were retained as the MP signature, consistent with prior work.^36^ MP activity was quantified in spatial transcriptomic data using gene set scoring, followed by per-observation standardization. Dominant MPs were assigned based on relative enrichment, and MP activity was summarized at the sample level and evaluated for associations with clinical features and cellular composition.

#### Gene set enrichment analysis of spatial molecular programs

Pathway enrichment analysis was performed on NMF-derived programs and metaprograms using gene sets from the Molecular Signatures Database (MSigDB; Hallmark and Gene Ontology collections). Program gene sets were compared to reference pathways using a one-sided hypergeometric over-representation test, with the background defined as the union of genes represented across all programs. Enrichment significance was computed as the upper-tail probability and transformed as – log_10_(p) for downstream analysis. Enrichment results were aggregated across programs to identify recurrent biological themes and visualized as clustered heatmaps of enrichment scores. For each enriched program–pathway pair, overlapping genes were recorded as lead genes to facilitate the interpretation of program-specific functional signatures.

#### Whorl–lobule trajectory analysis

Trajectory structure along the whorl–lobule axis was inferred using diffusion-based manifold learning. A random subset of cells was used to construct a low-dimensional embedding following imputation to denoise sparse transcriptomic measurements. Cells were embedded using PHATE, capturing continuous transcriptional structure associated with spatial molecular programs. Diffusion pseudotime was computed on the PHATE manifold to model progression along the whorl–lobule continuum. A neighborhood graph was constructed in the embedding space, followed by diffusion mapping and pseudotime inference. The root cell was defined as the cell with the lowest WL score, anchoring pseudotime to the WL state. To quantify directional structure along the manifold, endpoint populations were defined based on high expression of two spatial molecular programs (top 5% of program scores). Centroids of these populations were computed in PHATE space, and each cell was assigned a continuous score based on its relative distance to the two endpoints, providing a scalar measure of position along the whorl–lobule trajectory.

#### HIF-1*α* Immunohistochemistry

Monoclonal HIF-1*α* (EP1215Y; Biocare Medical) staining was performed at a 1:100 dilution. Heat-mediated antigen retrieval was performed using Tris/EDTA buffer at pH 9.0 for the staining. The secondary antibody used was MACH 2 Rabbit HRP-Polymer (RHRP520; Biocare Medical) with a 30-minute incubation period. Chromogen development was performed using Betazoid DAB buffer (DS900; Biocare Medical) with a 5-minute incubation period. The slides were then counterstained with hematoxylin Gill No. 3 (Avantik; RS4021).

### Statistical analyses

Statistical analyses were performed using two-sided tests unless otherwise specified. Correlations were assessed using Pearson or Spearman correlation coefficients as indicated. For comparisons between groups, appropriate non-parametric tests were used as indicated. P-values were adjusted for multiple comparisons using false discovery rate (FDR) correction where applicable.

#### Spatial Statistics

Spatial architecture and patterns of cell-type distribution were interrogated using Squidpy (v1.5.0). A spatial adjacency matrix was first constructed using sq.gr.spatial_neighbors(adata, coord_type=“generic“, spatial_key=“spatial“) to define neighboring cells based on Eu-clidean coordinates. Global spatial autocorrelation was quantified using Moran’s I via the sq.gr.spatial_autocorr function with 10 permutations to assess the clustering of specific cell types (e.g., Fibroblasts, T-cells) across different tumor grades. Neighborhood enrichment analysis was performed using sq.gr.nhood_enrichment to identify significant spatial interactions or exclusions between categorical groups, including CellType NanoString and dominant NMF components. Additionally, Ripley’s L-function, a linearized version of the K-function, was applied to 10% random subsamples of the data using sq.gr.ripley(mode=“L“) to characterize the scale of spatial clustering within clinical subgroups.

**Bray Curtis Dissimilarity** The Bray-Curtis dissimilarity is a metric commonly used in ecology to quantify the compositional dissimilarity between two different sites or samples. Mathematically, the Bray-Curtis dissimilarity between two landscapes, A and B, is defined as:

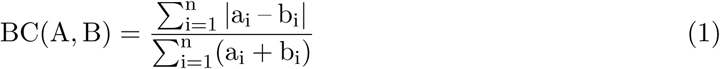

Where: n represents the total number of cell types, states, or ecotypes a_i_ and b_i_ are the counts of the i^th^ cell type or SMP in landscapes A and B.

In the context of our study on the ecological landscape of the meningioma microenvironment, each landscape corresponds to a specific combination of categories or metapopulations, and the cell types or SMPs represent distinct cellular populations or states within those landscapes. For each pair of landscapes, the Bray-Curtis dissimilarity quantifies the difference in their cellular composition. To assess the dissimilarity between landscapes, Bray-Curtis values were computed for each landscape, and mean cell-type values were extracted. The Bray-Curtis dissimilarity was then calculated for each pair of landscapes. The Bray-Curtis dissimilarity between all pairs of observations was calculated using the ‘beta_diversity‘ function from the ‘skbio.diversity‘ package.

#### Histology feature extraction from histology images

Histological features were quantified from nuclei and collagen image channels corresponding to Visium-profiled samples. Spatial coordinates were used to align gene expression measurements with histological images, and analyses were restricted to in-tissue spots with matched expression and imaging data. For each spot, fixed-size image patches (224 × 224 pixels) centered on spatial coordinates were extracted from nuclei and collagen channels. Images were converted to grayscale, intensity-normalized, and segmented using Otsu thresholding to identify signal-positive regions. Nuclei were segmented as discrete objects and quantified using object-based features, including count, density, and mean size, with size filtering applied to exclude debris and merged aggregates. Collagen content was quantified as the fraction of pixels classified as collagen-positive within each patch. Segmentation outputs were visually inspected across samples. All features were standardized within each sample using z-score normalization and integrated with spatial transcriptomic data for downstream analysis of tissue architecture and its relationship to SMPs.

### MeningNet prediction of spatial molecular programs from histology Histology patch extraction and preprocessing

Histology patches were extracted from two spatially resolved datasets: Visium whole-slide H&E images and NanoString TMA H&E images. For the Visium cohort, high-resolution histology images were paired with spatial coordinates from tissue_positions_list.csv files, and only spots annotated as in tissue were retained. For each retained Visium spot, a 256 × 256 pixel patch centered on the corresponding full-resolution coordinate was extracted. If a crop extended beyond image boundaries, the patch was zero-padded to preserve a fixed input size.

For the NanoString cohort, regions of interest were defined from ROI files corresponding to TMA fields of view (FOVs). Bounding boxes were extracted for each annotated FOV, optionally trimmed by a fixed margin, and subdivided into a regular grid; in the final training configuration, each FOV was divided into a 2 × 2 grid, yielding four image patches per FOV. Each patch was resized to 256 × 256 pixels before model input. Sample-specific orientation corrections were applied to selected NanoString images based on acquisition version metadata.

All image patches were stain-normalized using a custom LAB-space preprocessing routine in which the RGB image was converted to CIELAB color space, contrast-limited adaptive histogram equalization (CLAHE)^45^ was applied to the luminance channel, and the image was then converted back to RGB. Training images were augmented using TrivialAugmentWide,^46^ followed by tensor conversion and normalization to ImageNet mean and standard deviation (mean [0.485, 0.456, 0.406], standard deviation [0.229, 0.224, 0.225]). Validation and test images underwent the same stain normalization and ImageNet normalization without augmentation.

The final model was trained to predict eight retained spatial molecular targets, comprising six NMF-derived components (NMF1-NMF6) and two metaprogram scores (MP-9, MP-18). These targets were selected from a larger candidate set based on collinearity and clinical relevance.

#### MeningNet architecture and training criterion

We developed MeningNet, a histology-to-molecular prediction architecture that combines visual tokenization with transformer-based contextual modeling and output-specific attention pooling. Depending on the configuration, the model processes each input image either through non-overlapping patch embedding or through a pretrained convolutional stem based on ConvNeXt-Tiny.^47^ In our patch-based setting, an input image x ∈ R^H×W×3^ is partitioned into non-overlapping patches of size P × P, flattened, linearly projected into a d-dimensional embedding space, and normalized. In the convolutional setting, the image is first passed through the ConvNeXt-Tiny feature extractor, and the resulting feature map is projected into the same transformer embedding dimension using a 1D convolution. In both cases, the spatial features are reshaped into a sequence of visual tokens and combined with learned positional embeddings before transformer encoding.

### Specifically, let X^(0)^ = {x_1_, *. . .*, x_N_}, x_i_ ∈ R^d^, denote the tokenized image representation, where N is the number of spatial tokens. Learned positional embeddings E are then added to obtain

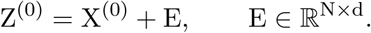

Motivated by advances in vision transformers,^48^ then the token sequence is processed by a stack of transformer encoder blocks composed of pre-normalized multi-head self-attention and feedforward sublayers with residual connections:

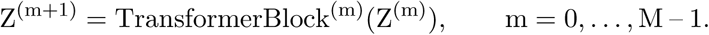

The final contextualized representation is Z ∈ R^N×d^.

Rather than using a dedicated class token, MeningNet employs learned output-specific attention pooling to derive one latent representation per molecular component. Specifically, the model computes attention logits over spatial tokens for each output component, normalizes them across token positions with a softmax, and uses the resulting weights to form a weighted sum of the contextualized token features. This yields a set of component-specific embeddings

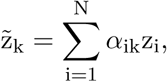

where *α*_ik_ denotes the attention weight assigned to token i for component k. Each component embedding ẑ_k_ is then passed through its own prediction head, consisting of layer normalization, a hidden linear projection, GELU activation, dropout, and a final linear layer, to produce the corresponding scalar output. This design allows the model to learn spatially distinct evidence patterns for different molecular programs while preserving a shared image encoder.

Our MeningNet also supports optional task-level auxiliary supervision. When auxiliary tasks are defined, a parallel attention-pooling branch computes task-specific pooled representations from the same transformer tokens. These task representations are then passed through dedicated task heads to generate task-level logits. Optionally, the mean component representation can be fused with the task representation prior to classification, thereby enabling controlled information sharing between the primary regression outputs and auxiliary tasks.

**Component-specific attention pooling** To generate predictions for the K molecular components, MeningNet aggregates the contextualized token representations using a learned component-specific attention pooling mechanism. Given the final transformer output Z = {z_1_, …, z_N_}, z_i_ ∈ R^d^, the model first computes attention logits for each token-component pair, H = g_comp_(Z) ∈ _R_N×K.

These logits are then normalized across spatial tokens for each component using a softmax,

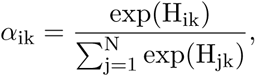

where *α*_ik_ denotes the contribution of token i to molecular component k. The resulting attention weights are used to form component-specific latent representations,

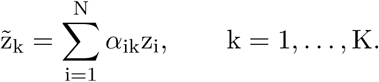

Each pooled representation ẑ_k_ is then passed through an independent prediction head h_k_(·) to produce a scalar output,

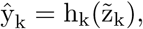

and the final molecular prediction vector is

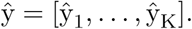

This output-specific pooling strategy allows the model to identify distinct spatial evidence patterns for different molecular programs while maintaining a shared transformer backbone.

**Auxiliary clinical prediction tasks** In addition to molecular regression, MeningNet supports auxiliary clinical prediction tasks through a parallel task-specific attention branch applied to the same contextualized token sequence Z = {z_1_, *. . .*, z_N_}, where z_i_ ∈ R^d^. For each auxiliary task t, the model computes task-specific attention logits over the N spatial tokens, followed by softmax normalization to obtain attention weights *α*^task^. These weights are used to form a task-specific pooled representation

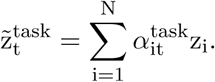

To incorporate molecular context into the auxiliary predictions, the model computes the mean component representation

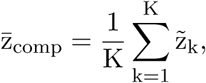

and, when enabled, adds this shared molecular summary to the task-specific representation before classification:

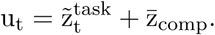

The fused representation u_t_ is then passed through a task-specific prediction head to produce the corresponding logits,

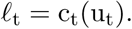

This design allows the auxiliary clinical predictions to be informed both by task-relevant spatial patterns and by the global molecular representation learned by the model. During training, auxiliary classification loss is computed only for samples with valid task labels, and samples lacking annotations for a given task are masked from that task-specific loss term.

**Loss function.** We trained our MeningNet with a composite objective designed to balance accurate regression with preservation of the relative molecular structure across outputs. Because molecular programs are often most informative not only in their absolute magnitude but also in their pattern across components, we combined a Smooth L1 regression term with Pearson-correlation-based regularization at both the full-vector and component levels. Let y ∈ R^K^ denote the ground-truth molecular target vector and y^ ∈ R^K^ the corresponding prediction. The main regression objective is

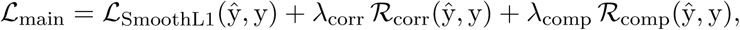

where R_corr_ encourages agreement between the predicted and observed molecular profiles, and R_comp_ further regularizes component-wise consistency. This formulation encourages the model to reduce absolute prediction error while preserving the relative organization of the molecular program.

When auxiliary supervision was available, we added task-specific cross-entropy losses to support joint learning of molecular and clinical targets:

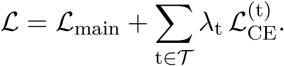

Here, T denotes the set of auxiliary tasks and *λ*_t_ the corresponding task weight. In practice, regression loss was computed only on samples with valid molecular targets, and missing targets were excluded from loss evaluation. During MixUp or CutMix augmentation, the regression objective was computed using the mixed targets, whereas auxiliary classification loss was evaluated on the corresponding clean samples.

#### Implementation and training details

All models were implemented in PyTorch and trained on A100-GPU when available. For the final training recipe, we used the hybrid transformer variant with a pretrained convolutional stem, instantiated with input size 256 × 256, patch size 16, embedding dimension 1024, transformer depth 10, 16 attention heads, multilayer perceptron dimension 1024, and prediction-head hidden dimension 1024. Dropout and embedding dropout were both set to 0.4, stochastic depth was set to 0.4, and attention dropout was set to 0.3.

Data were partitioned at the patient level using YaleID to prevent cross-patient leakage. When available, predefined patient split files were used; otherwise, grouped random splitting was performed using a 70%/15%/15% train/validation/test ratio. Training used the combined dataset configuration with NanoString upsampling factor 4 and grid size (2, 2). The optimization objective was a regression-correlation loss with a Smooth L1 base term and Pearson-correlation regularization, with a correlation weight of 0.4 and a component-level correlation weight of 0.2. For optimization, we used AdamW^49^ with weight decay 0.01. During the final training run, the ConvNeXt-tiny stem was partially fine-tuned: all backbone parameters were frozen except for the last three stages, and discriminative learning rates were applied across parameter groups. The transformer and non-backbone parameters used a learning rate of 1 × 10^−5^, whereas ConvNeXt stages 7, 6, and 5 used learning rates scaled by 0.1, 0.05, and 0.005, respectively. The learning rate was scheduled using cosine annealing with warm restarts (T_0_ = 8, T_mult_ = 2, *η*_min_ = 10^−8^).

Training was performed for up to 1000 epochs with early stopping based on validation loss and a patience of 15 epochs. At each optimization step, gradients were clipped to a maximum norm of 1.0. The best-performing model state on the validation set was checkpointed and reloaded at the end of training. Training artifacts, including checkpoints, loss curves, prediction plots, and attention visualizations, were written to structured output directories.

With probability 0.9, each batch underwent either CutMix^50^ or MixUp;^51^ CutMix was selected with probability 0.5, and both augmentations used *α* = 1.0. Regression loss for augmented batches was computed using the mixed targets. When auxiliary task supervision was enabled, task-classification loss was computed on the corresponding clean batch and added to the regression objective. Samples with missing regression targets were excluded from the regression loss computation using a validity mask based on NaN detection.

Model selection and evaluation were based on validation loss. During validation and test-time evaluation, we computed loss, cosine similarity, overall Spearman correlation, and overall R^2^, as well as per-component Spearman correlation and per-component R^2^. When auxiliary tasks were present, task-specific classification accuracies were also tracked. For the best validation epoch, regression scatter plots and component-attention heatmaps were additionally saved for qualitative inspection.

#### Whole-Slide Aggregation and Clinical Outcome Evaluation

**Patch-level prediction aggregation.** Patch-level prediction scores generated from whole-slide inference were stored as Apache Parquet files, each containing per-tile model outputs and

spatial coordinates. For each slide, component-wise prediction columns were extracted and aggregated to derive slide-level summary statistics.

For each prediction target, global mean predicted scores were computed across all tiles:

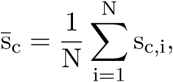

where s_c,i_ denotes the predicted probability for component c at tile i, and N is the total number of tiles.

To capture spatially localized high-confidence regions, hotspot-based aggregation was performed by ranking tile-level predictions and computing the mean of the top k% values:

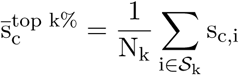

where S_k_ is the set of indices of the top k% tiles ranked by s_c_.

**Spatial composition analysis.** For each tile, the dominant predicted component was defined as the index of the maximum predicted probability across all components:

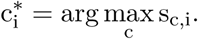

Slide-level spatial composition was then quantified as the proportion of tiles assigned to each component:

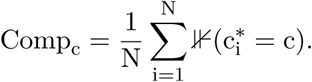

**Binary outcome definition.** Clinical endpoints were binarized for classification tasks. Tumor grade was encoded as high-grade (CNS WHO grade 2/3) versus low-grade (CNS WHO grade 1). Recurrence status was defined as a binary indicator based on clinical annotations (e.g., yes/no, 1/0). Model performance was evaluated using receiver operating characteristic (ROC) analysis. For each aggregation strategy (mean and top 10%), area under the ROC curve (AUC) was computed. An optimal classification threshold was selected using the Youden index and corresponding sensitivity and specificity were computed from the confusion matrix. To assess statistical robustness, performance metrics (AUC, sensitivity, specificity) were estimated using stratified bootstrap resampling (1,000 iterations) at the patient ID level. For each iteration, samples were drawn with replacement while preserving class balance, and metrics were recomputed. Confidence intervals were defined as the 2.5th and 97.5th percentiles of the bootstrap distributions.

#### Post Hoc Error-Stratified and Component-Level Analyses

Because the upstream evaluation output encoded correct predictions as a single category (Correct) and incorrect predictions as FP or FN, true positives (TP) and true negatives (TN) were reconstructed post hoc using the binary ground-truth label. This was applied separately for grade-mean, grade-top10, recurrence-mean, and recurrence-top10 prediction outputs, yielding four confusion-stratified categorical variables.

For each error-stratified grouping, descriptive statistics were computed for total tile count and slide-level prediction scores. Probability distributions and total tile counts were visualized by confusion stratum. To identify features associated with specific error modes, pairwise non-parametric comparisons were performed using two-sided Mann–Whitney U tests for FP versus TN and FN versus TP contrasts. Effect sizes were quantified using Cliff’s delta, and resulting P-values were adjusted for multiple testing using the Benjamini–Hochberg false discovery rate procedure.

**Dominant-component assignment and comparison.** At the patient level, the dominant component was defined as the component with the maximum component fraction across the renamed component columns:

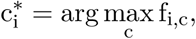

where f_i,c_ is the normalized fraction of component c in patient i. Component composition across confusion strata was summarized by averaging component fractions within each stratum and by quantifying the proportion of cases for which each component was dominant. Associations between categorical dominant-phenotype labels and confusion strata were assessed using contingency tables and chi-square tests where noted.

**Diagnosis-level component enrichment and dominant-component prevalence.** For diagnosis-level analyses, component fractions were first normalized within each patient:

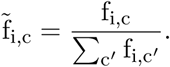

Within each diagnosis group, the mean component fraction was computed for each component, and component enrichment was expressed as a z-score relative to the diagnosis-restricted cohort:

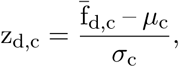

where f_d,c_ is the mean fraction of component c among cases with diagnosis d, and *µ*_c_ and *σ*_c_ are the cohort-wide mean and standard deviation of that component. Dominant-component prevalence was defined as the percentage of cases within a diagnosis for which component c was the dominant component:

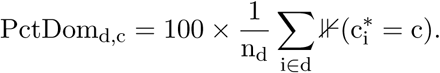

## Resource Availability

### Data Availability

Data generated in this study have been deposited in the NCBI Gene Expression Omnibus (GEO) and are currently held private pending journal publication. The dataset comprises a spatially resolved, multi-omic atlas of human meningioma with matched histology, including single-cell RNA sequencing (220,043 cells), 10x Visium spatial transcriptomics (56,020 spots across 17 sections), NanoString CosMx RNA spatial transcriptomics (391,051 cells across 77 tumors), and NanoString CosMx Protein spatial proteomics (993,675 cells across 119 tumors). Ten scRNA-seq samples included in the atlas were obtained from a previously published study^19^ and are available through SRA BioProject PRJNA660307. External validation data from the Digital Brain Tumor Atlas were obtained from publicly available sources, as described in the original studies.^38^

## Code Availability

All code used for data processing, spatial analysis, and model development is available at https://github.com/dmiyagi/meningioma_spatial. The repository includes pipelines for preprocessing and integrating scRNA-seq, Visium, and CosMx datasets; deriving spatial molecular programs using sample-wise non-negative matrix factorization and consensus metaprogram analysis; and performing spatial statistical analyses, along with tutorials for all major figures. Repository used for training a deep learning model, as well as the downstream scripts used for evaluation, and sliding-window WSI inference pipelines are provided, together with pretrained model weights and example notebooks. Any additional information required to reanalyze the data reported in this paper is available from the lead contact upon request.

## Author contributions

D.F.M. and M.G. conceived the study. D.F.M. and A.A. developed the methodology. D.F.M., D.M., E.Z.E-O., K.Yal., Y.T., P.Q.D., B.G., J.Y., K.M.-G., K.Yas., G.W., A.G.E.-S., J.M., and T.B. performed critical aspects of the investigation. D.F.M., M.W.Y., J.Y., K.M.-G., and J.M. curated the data. D.F.M., A.A., and E.Z.E-O. performed formal analyses. G.W., J.Y., N.S., J.M., T.B., and M.G. provided resources. D.F.M. wrote the original draft with input from A.A., T.B., D.M. and M.G. D.F.M., A.A., D.M., E.Z.E-O., R.G.W.V., J.M., T.B., and M.G. reviewed and edited the manuscript. M.G. supervised the study.

## Acknowledgments

This work was supported by the Connecticut Brain Tumor Alliance and NIH grant 1R01NS110824-01A1 (for work with TRAF7-mutant meningiomas only) to M.G. D.F.M. was supported by NIH/NCI (no. F31CA254426) and NIH–Medical Scientist Training Program (no. T32GM007205). We would like to acknowledge Kaya Bilguvar, MD, PhD, Chris Castaldi, Amos Brooks, Lori Charette, and Yalai Bai, MD, PhD, for technical assistance at the Yale Center for Genome Analysis and Yale Pathology tissue services.

## Declaration of generative AI and AI-assisted technologies in the manuscript preparation process

During the preparation of this work, the authors used large language models Codex and Gemini for limited support in code development, and Grammarly for manuscript editing. All computational analyses, model development, and biological interpretations were conceived, implemented, and verified by the authors. After using these tools, the authors reviewed and edited the content as needed and take full responsibility for the content of the published article.

---

**Figure S1.**
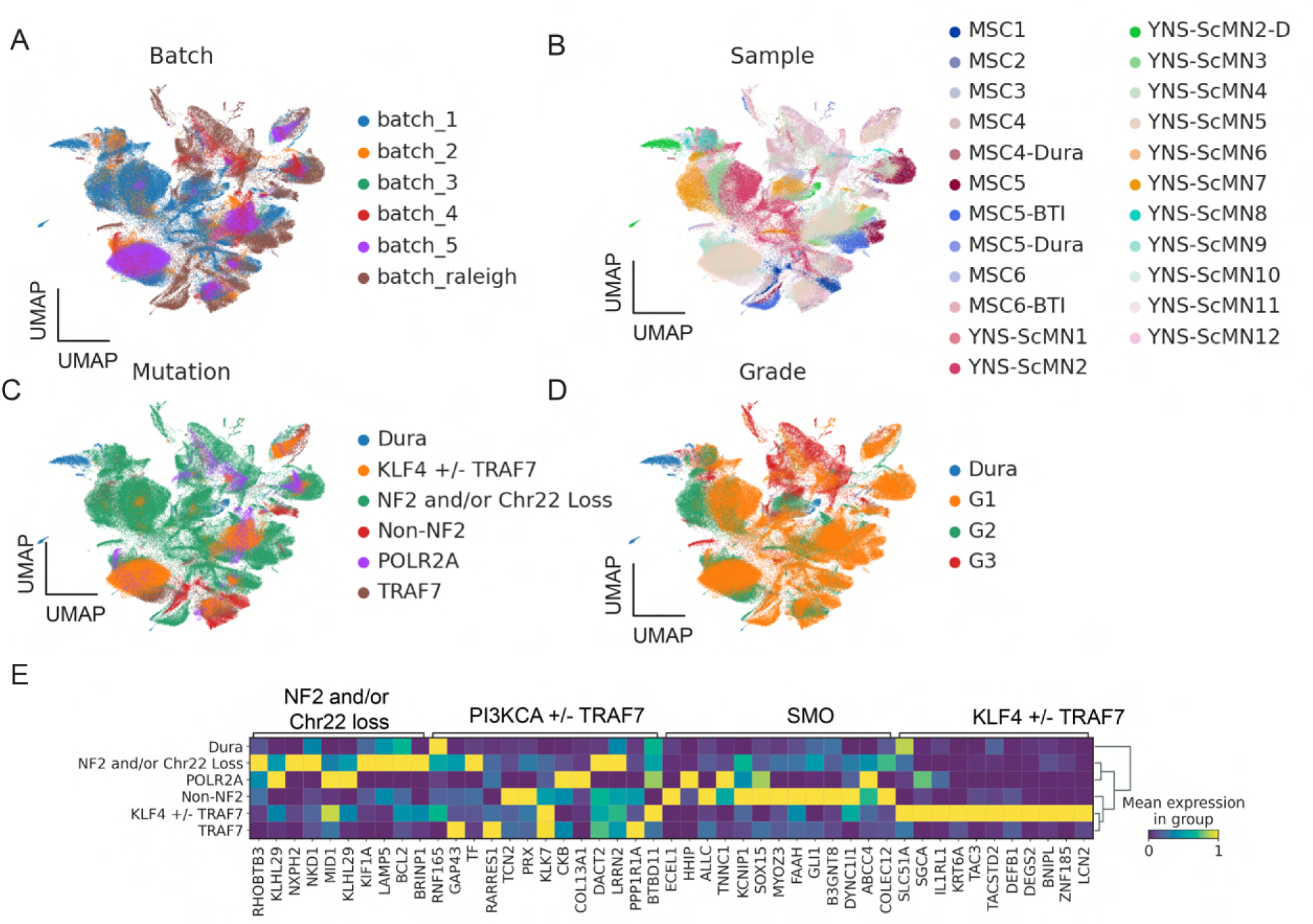
Single Cell Sequencing Batch Correction and Expression Comparison. Principal Component Analysis (PCA) visualizes 20 meningioma samples and 3 dura samples included in this study. **A.** PCA of batch correction, **(B)** sample distribution, **(C)** mutation profiles, **(D)** and tumor grades. **E.** Aggregated sc-RNAseq samples by genetic profile *(left)* and compared with the top 10 genes previously identified as specific to genomic subgroups *(top)*.^11^

**Figure S2.**
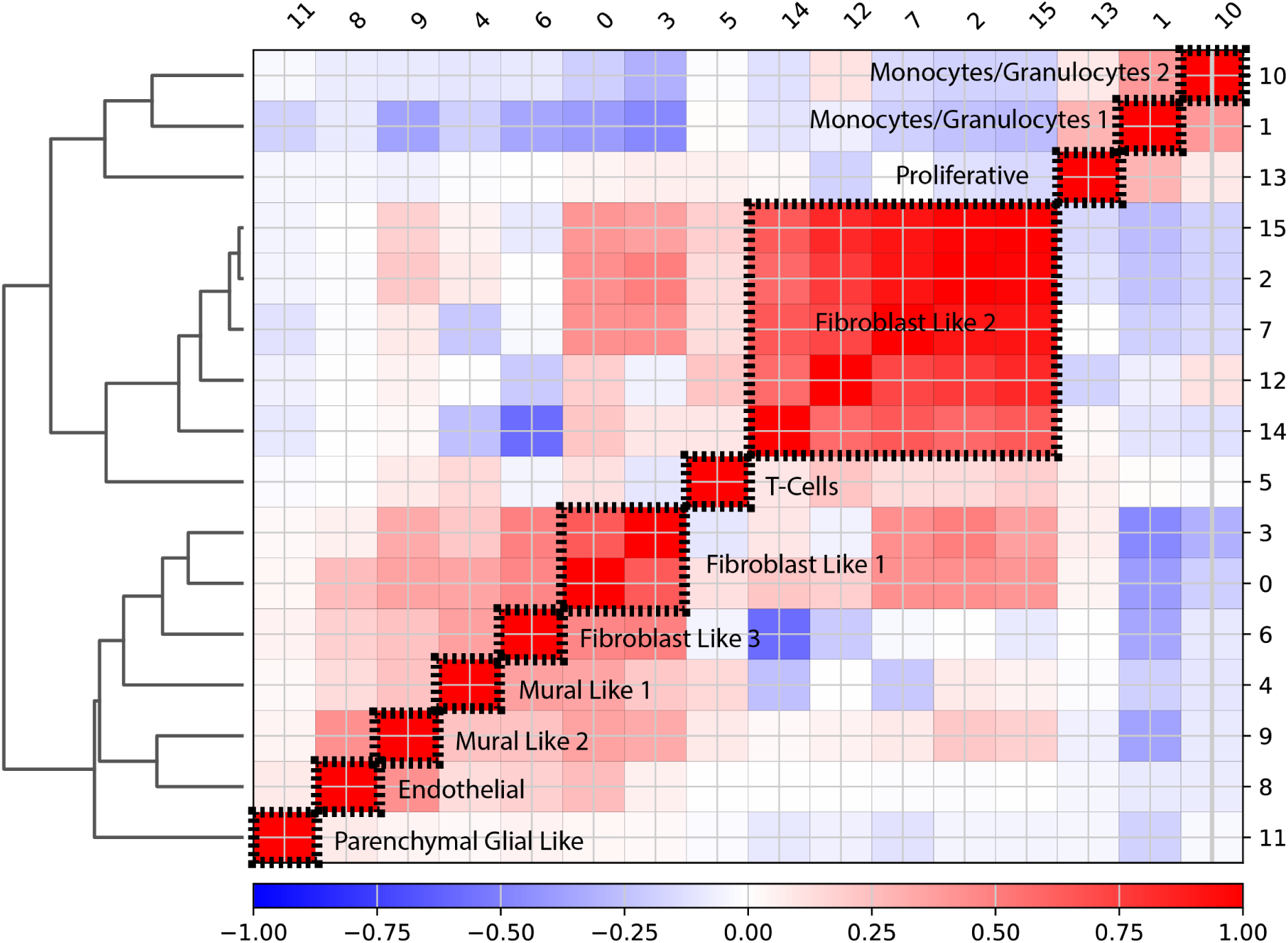
Correlation Matrix of Annotated Unsupervised sc-RNA Clusters. The correlation matrix depicts the relationships between annotated unsupervised single-cell RNA (sc-RNA) clusters. The sample dendrogram is shown on the *left*, representing the hierarchical unsupervised clustering. The Pearson correlation *r* values are displayed along the *bottom* of the matrix, ranging from −1 to +1, indicating the strength and direction of correlations. The original unsupervised cluster numbers are indicated on the *right* and *top* sides of the matrix. A threshold of Pearson *r* > 0.5 was used to determine cluster amalgamation.

**Figure S3.**
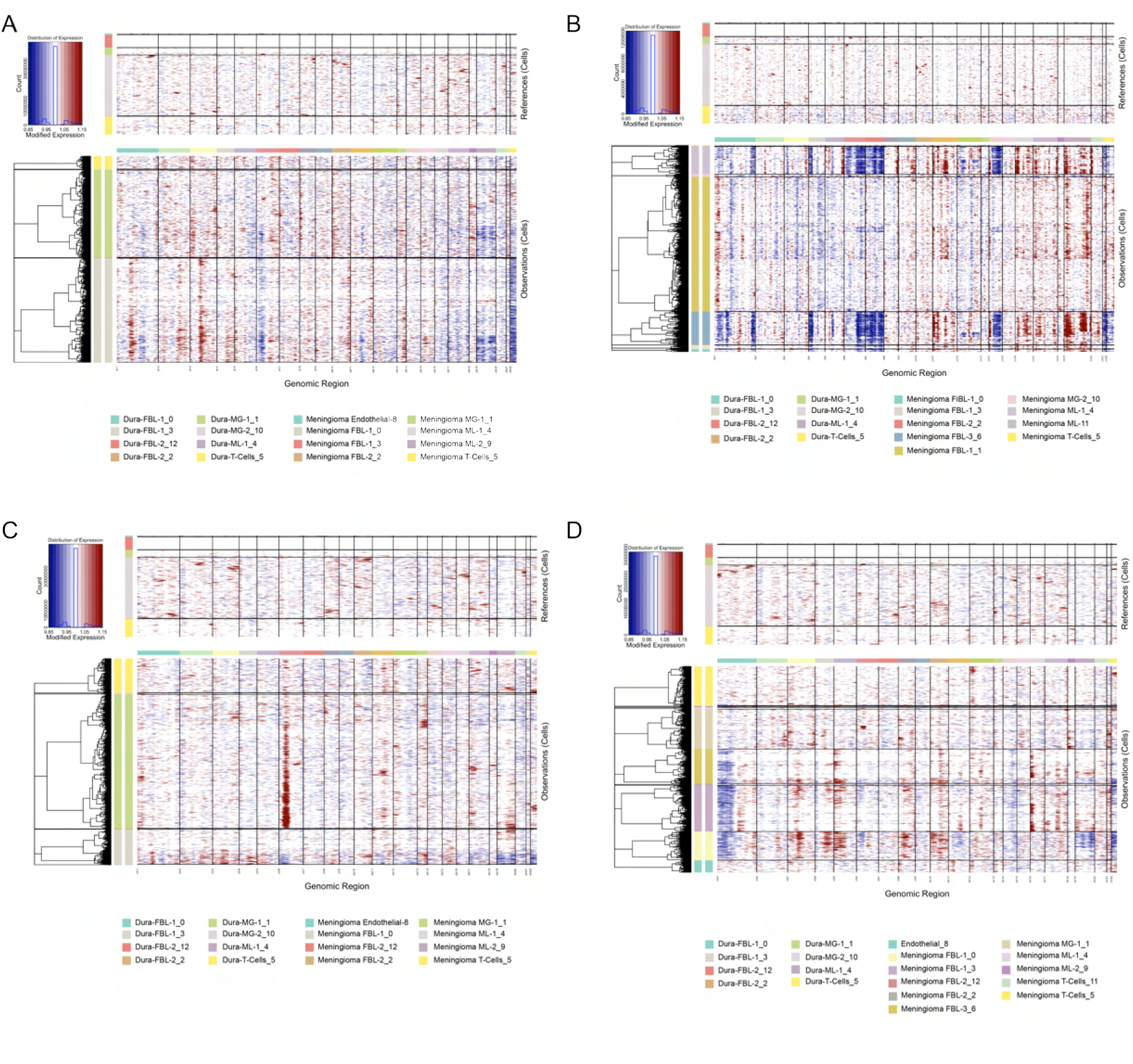
InferCNV heatmaps demonstrating chromosome events in four samples used to annotate chr22q/chr1p loss. **A.** Sample YNS-scMN3 with chr22q loss. **B.** Sample YNS-scMN8 with chr1p/chr22q loss. **C.** Sample YNS-scMN7 with chr22q loss. **D.** Sample YNS-scMN2 with chr1p/chr22q loss.

**Figure S4.**
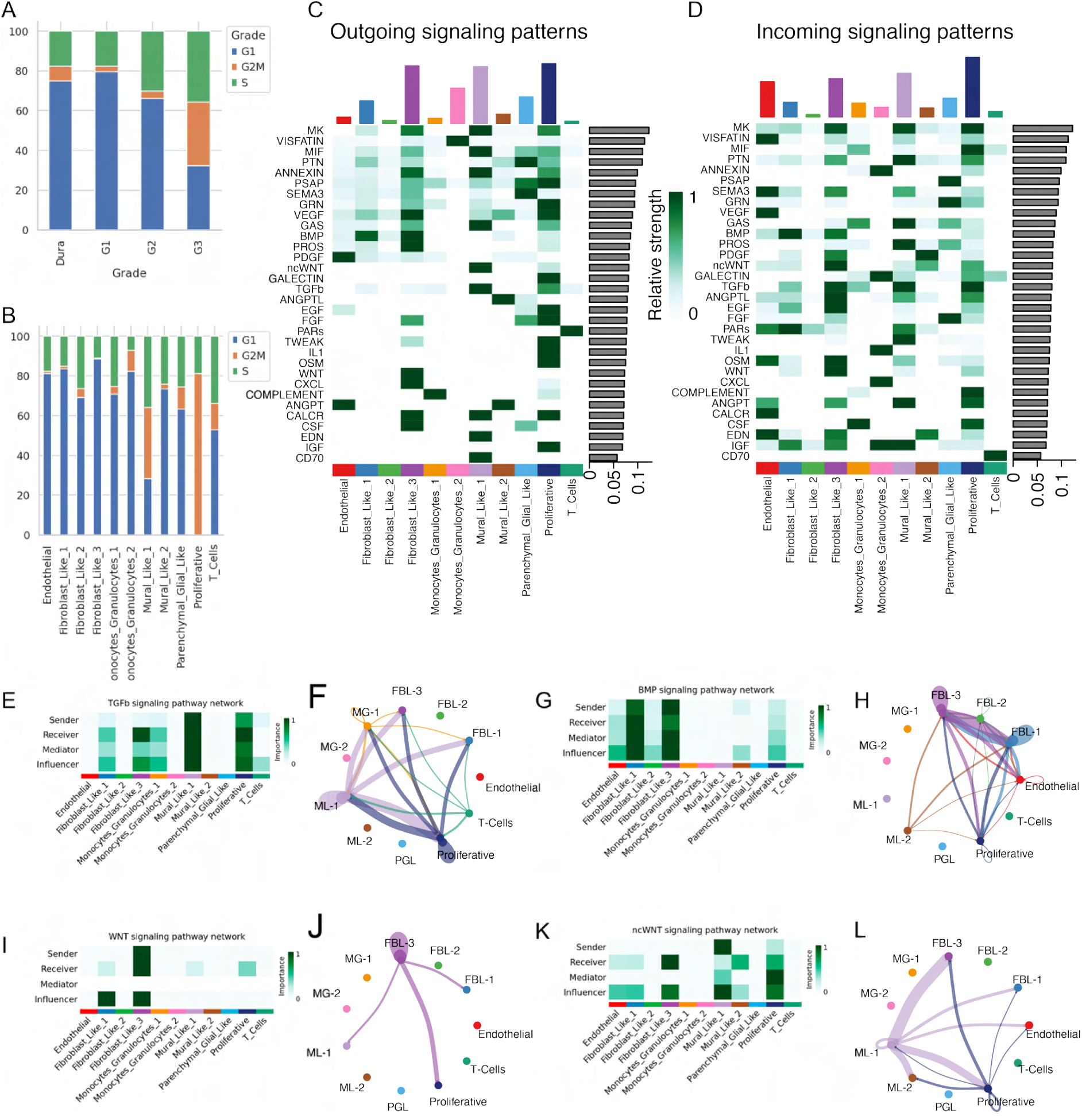
Identification of malignant cell populations and intercellular signaling networks. **A.** Cell cycle state distribution across samples stratified by tumor grade, based on canonical G1/S/G2M gene signatures.^28^ Bars represent the proportion of cells assigned to each phase. **B.** Cell cycle state distribution across scRNA-defined cell types, highlighting enrichment of proliferative states within specific populations. **C–D.** Cell–cell communication inferred using CellChat. Heatmaps show relative outgoing (C) and incoming (D) signaling strengths for major ligand–receptor pathways across cell types. Rows represent signaling pathways, columns represent cell types, and color intensity indicates relative signaling strength. *Continued on next page*. **E, G, I, K.** Pathway-specific signaling role analysis for TGF*β*, BMP, WNT, and non-canonical WNT (ncWNT). Heatmaps depict the relative contribution of each cell type as sender, receiver, mediator, and influencer within each signaling pathway. **F, H, J, L.** Corresponding network visualizations of pathway-specific interactions, illustrating the directionality and strength of inferred signaling between cell types. Node identity denotes cell type, and edge thickness reflects interaction strength.

**Figure S5.**
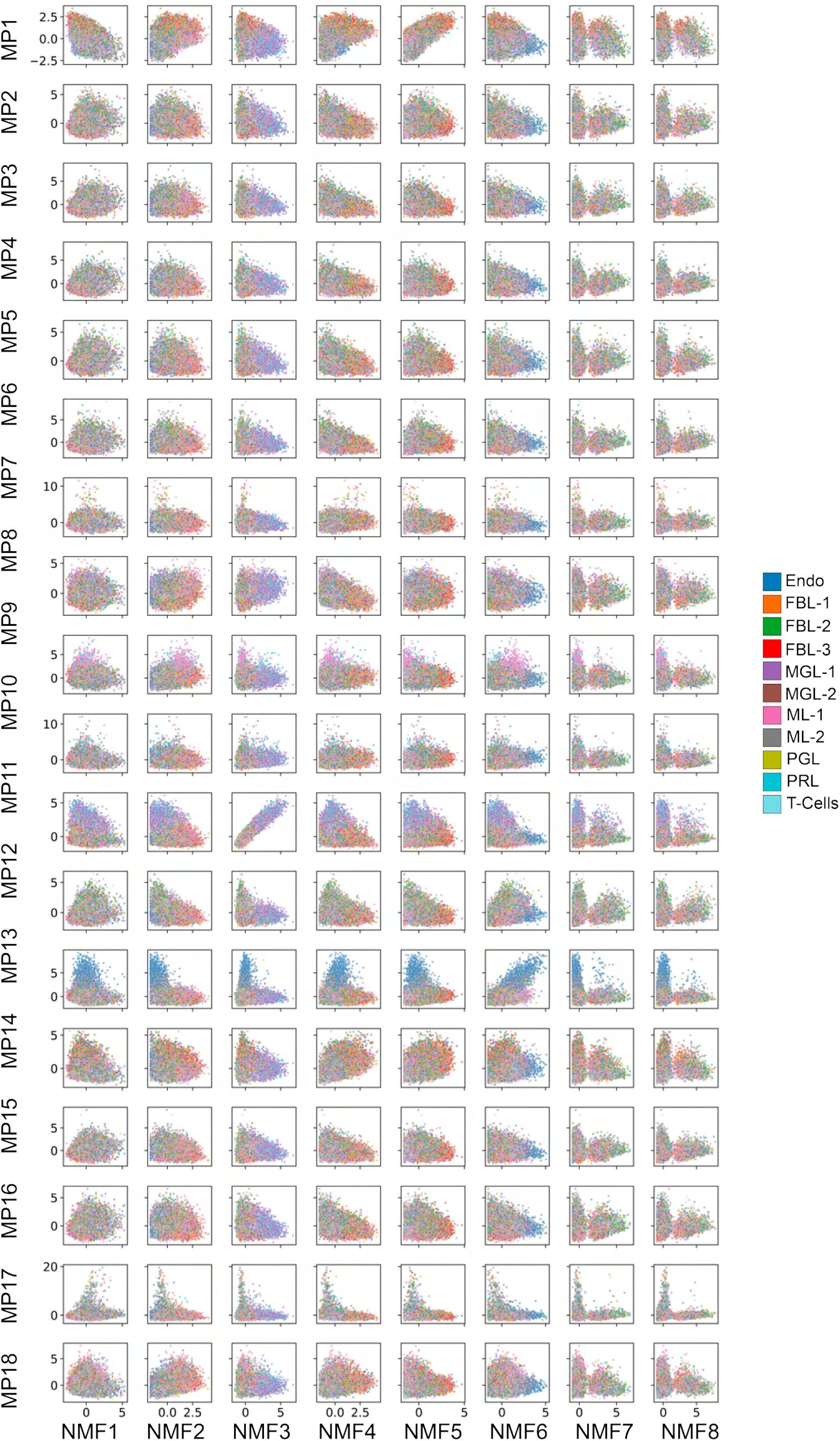
Relationship between RNA meta-programs (MPs) and NMF components across single cells. Scatter plots showing per-cell scores for each meta-program (MP1–MP18; rows) versus each RNA-NMF component (NMF1–NMF8; columns). Each point represents a single cell, colored by scRNA-defined cell type. A random subset of 10% of cells is shown for visualization. Quantitative correlation values for all MP–NMF pairs are provided in Supplementary File 5.

**Figure S6.**
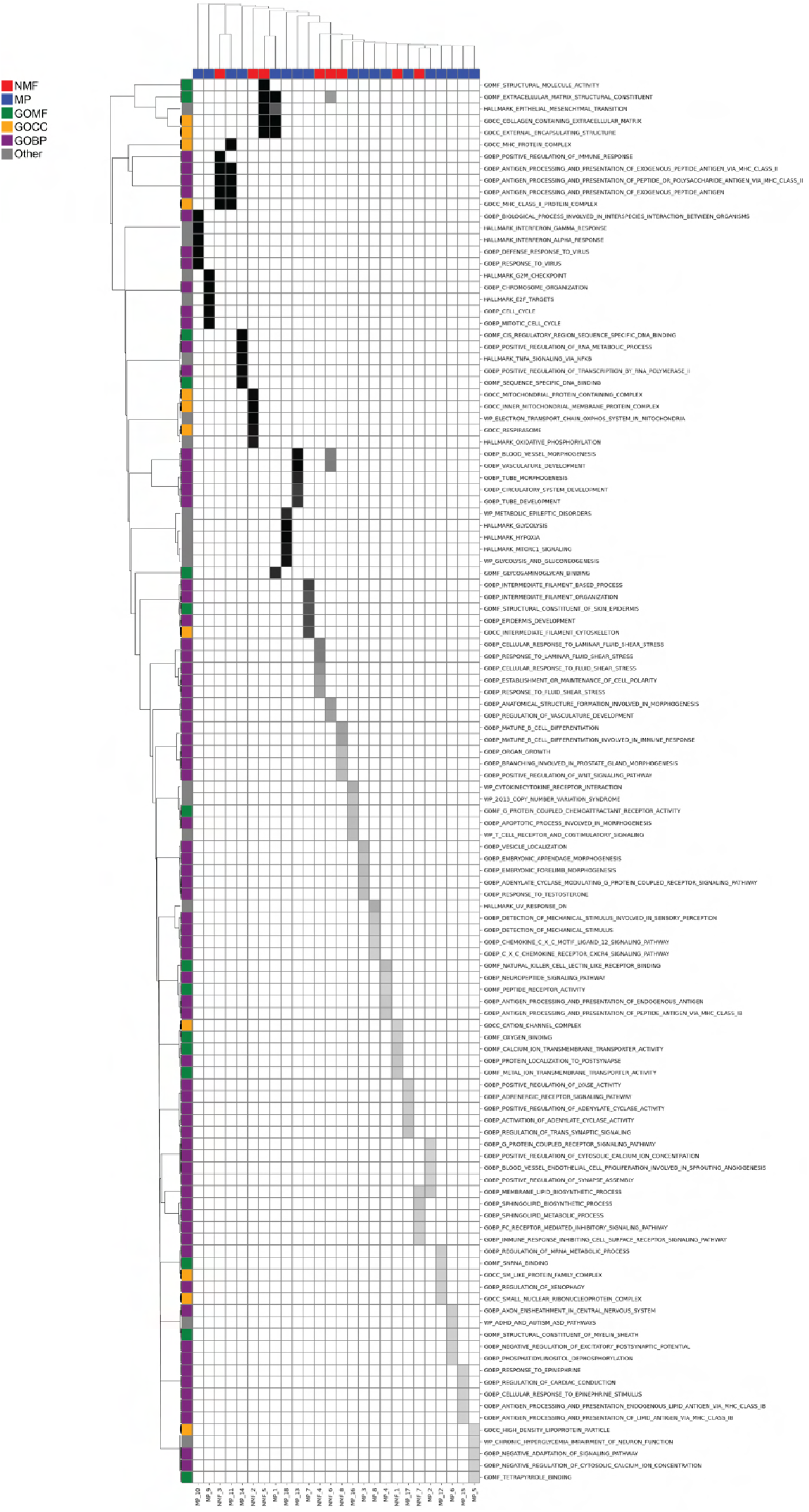
Spatial Network Pathway Enrichment. Rows correspond to gene programs: GO Molecular Functions (GOMF), GO Cellular Components (GOCC), GO Biological Processes (GOBP), Other corresponds to HallmarkDB or WikiPathways as denoted by gene program prefix. Columns correspond to SMPs, NMF (red), MP (blue).

**Figure S7.**
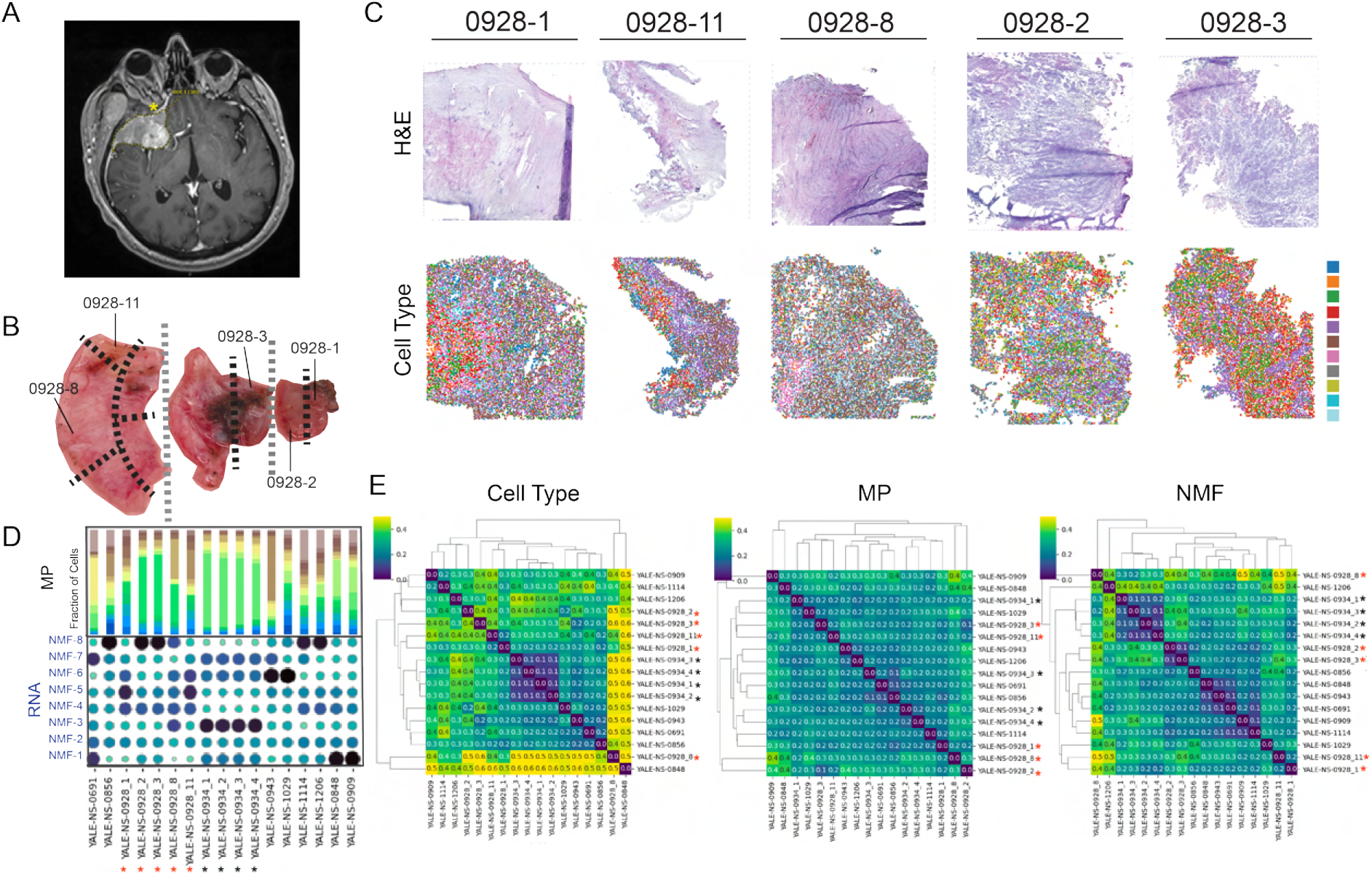
Multi-region spatial profiling of Meningiomas using Visium . **A.** T1-weighted post-contrast MRI showing a right sphenoid wing meningioma, with hyperostosis and dural thickening (arrow). **B.** Gross pathology and sampling schematic for a WHO grade 1 KLF4/TRAF7 meningioma. Five regions (core and tumor-dura interface) were sampled for spatial transcriptomics. **C.** H&E images (top) and spatial cell-type maps (bottom) for the five sampled tumor sections from patient 0928. **D.** MP scores (top bar) overlaid on RNA-NMF usage (dot plot) across spatial sections. **E.** Bray–Curtis dissimilarity matrices of spatial sections, based on cell type (left), MP (middle), and NMF (right). Stars mark sections from the same tumor. MP and NMF-based clustering yielded lower intra-tumor dissimilarity than cell-type-based annotation. Clustering based on RNA-defined cell types, protein-defined cell types, MP (RNA), NMF (RNA), and NMF (protein) programs. Histology annotations (e.g., anaplastic, secretory, transitional) are shown.

**Figure S8.**
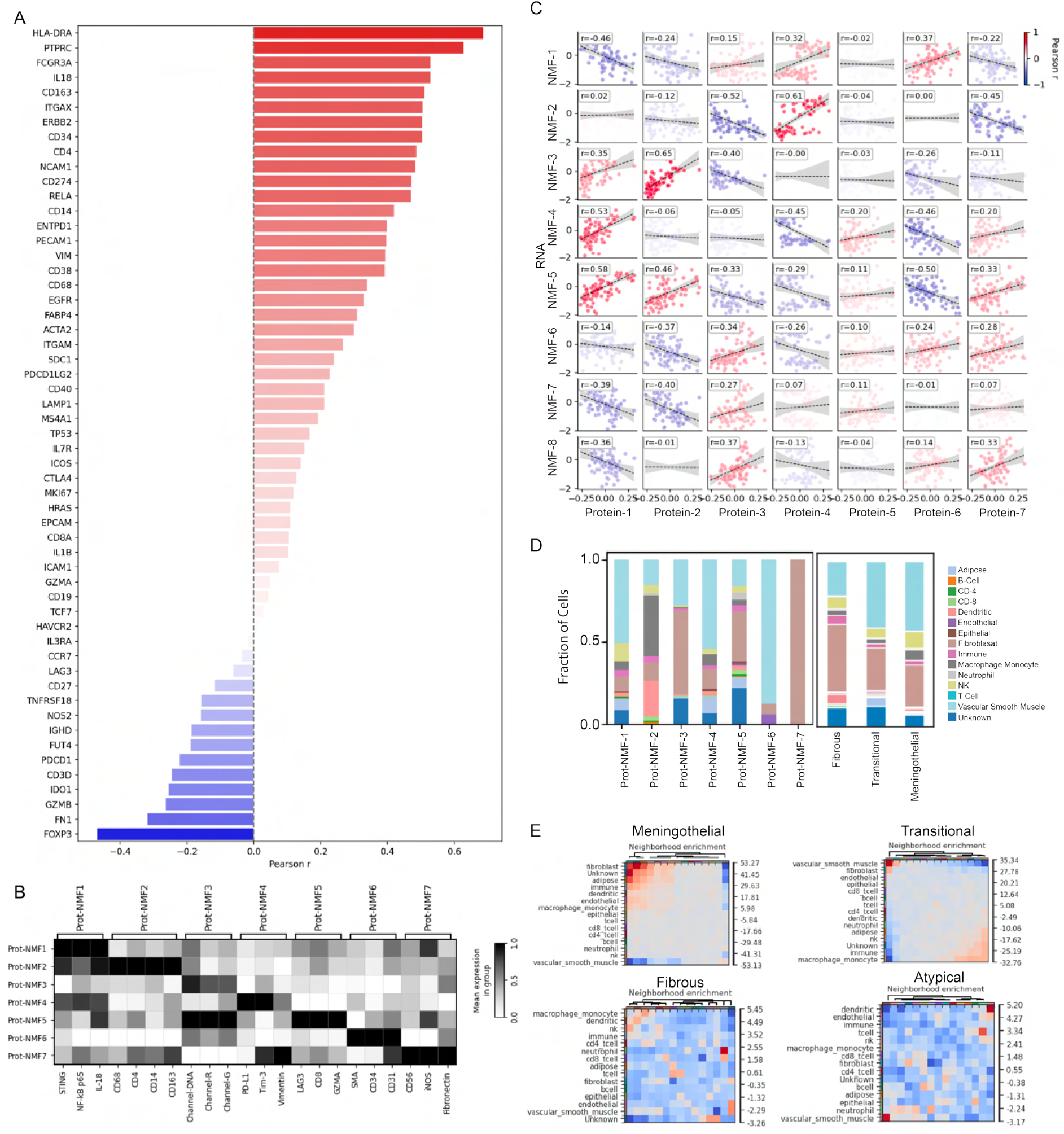
NanoString CosMx spatial proteomic analysis and integration with RNA-derived programs. **A.** Gene-level correspondence between RNA and protein modalities. Bar plot showing Pearson correlation coefficients between matched RNA transcripts and protein expression across samples, highlighting concordant and discordant markers. **B.** Protein-derived NMF components. Heatmap of scaled protein expression across NMF-derived protein components (Protein-NMF 1–7), annotated by representative marker proteins. **C.** Cross-modality program alignment. Pairwise Pearson correlations between RNA-derived NMF components and protein-derived NMF components across matched regions, indicating correspondence between transcriptional and proteomic programs. **D.** Cellular composition of protein-defined programs. Stacked bar plots showing the fraction of NanoString-defined cell types within each protein NMF component (left), and across histologic subtypes, including fibrous (n=3), transitional (n=20), and meningothelial (n = 30) (right), illustrating multicellular composition of proteomic programs. **E.** Spatial neighborhood enrichment analysis. Heatmaps showing enrichment or depletion of neighboring cell types within histologic subtypes, including meningothelial, transitional, fibrous, and atypical (n = 28), highlighting distinct microenvironmental architectures associated with each histology.

**Figure S9.**
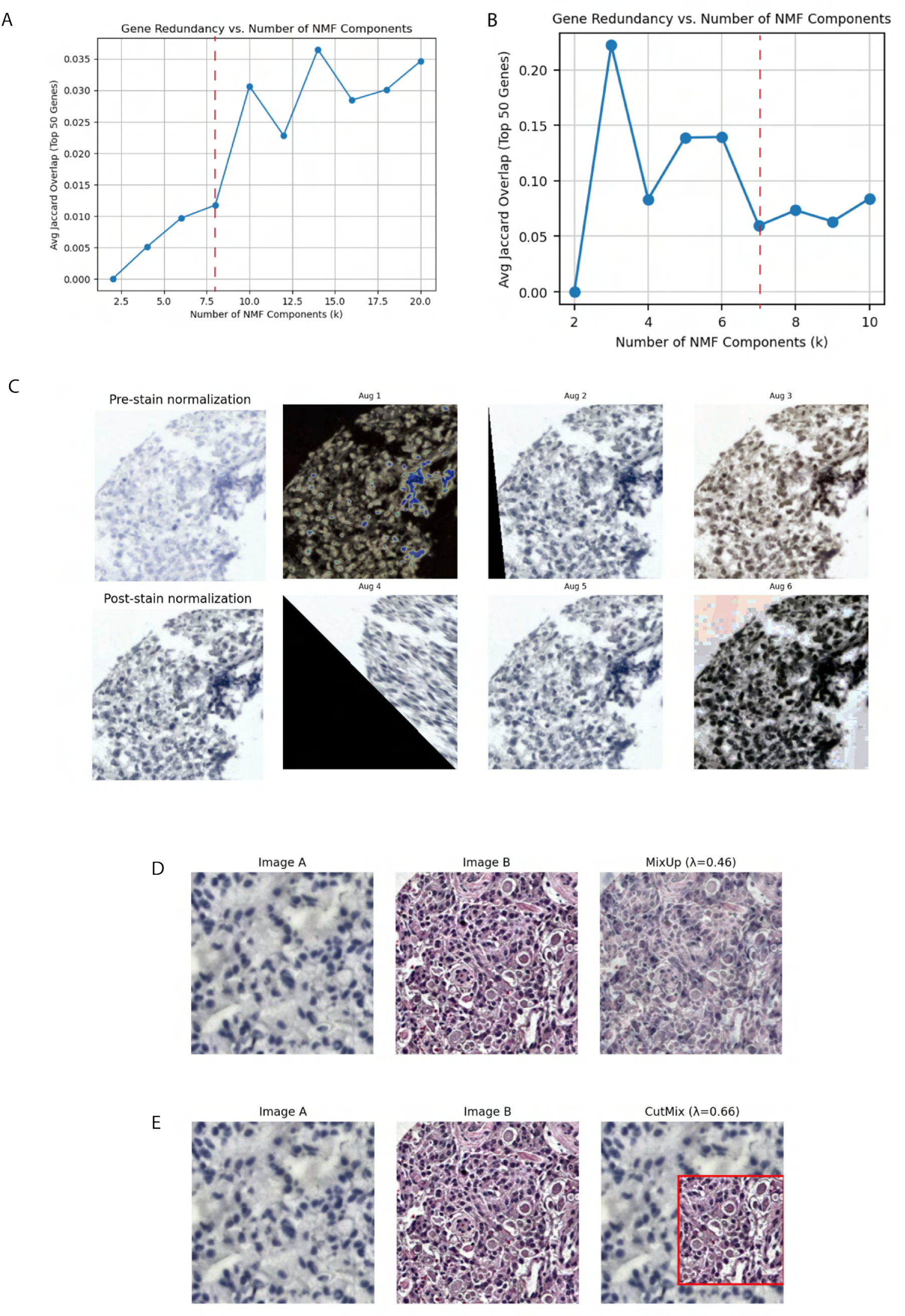
Selection of NMF rank and image preprocessing and augmentation strategies. **A–B.** Selection of optimal factorization rank for non-negative matrix factorization (NMF) based on gene redundancy. Average Jaccard overlap of top-ranked genes is plotted as a function of the number of components (k) for RNA (**A**) and protein (**B**) datasets. Dashed red lines indicate the selected rank corresponding to the first local minimum in inter-component overlap, marking the onset of increasing redundancy. **C.** Representative H&E image patches before and after stain normalization, illustrating reduction in color variability across samples. Example augmented patches are shown, demonstrating transformations applied during training. **D.** MixUp augmentation, in which two input images are linearly combined to generate a synthetic training example. **E.** CutMix augmentation, in which a region from one image is replaced with a patch from another image to enhance model robustness.

**Figure S10.**
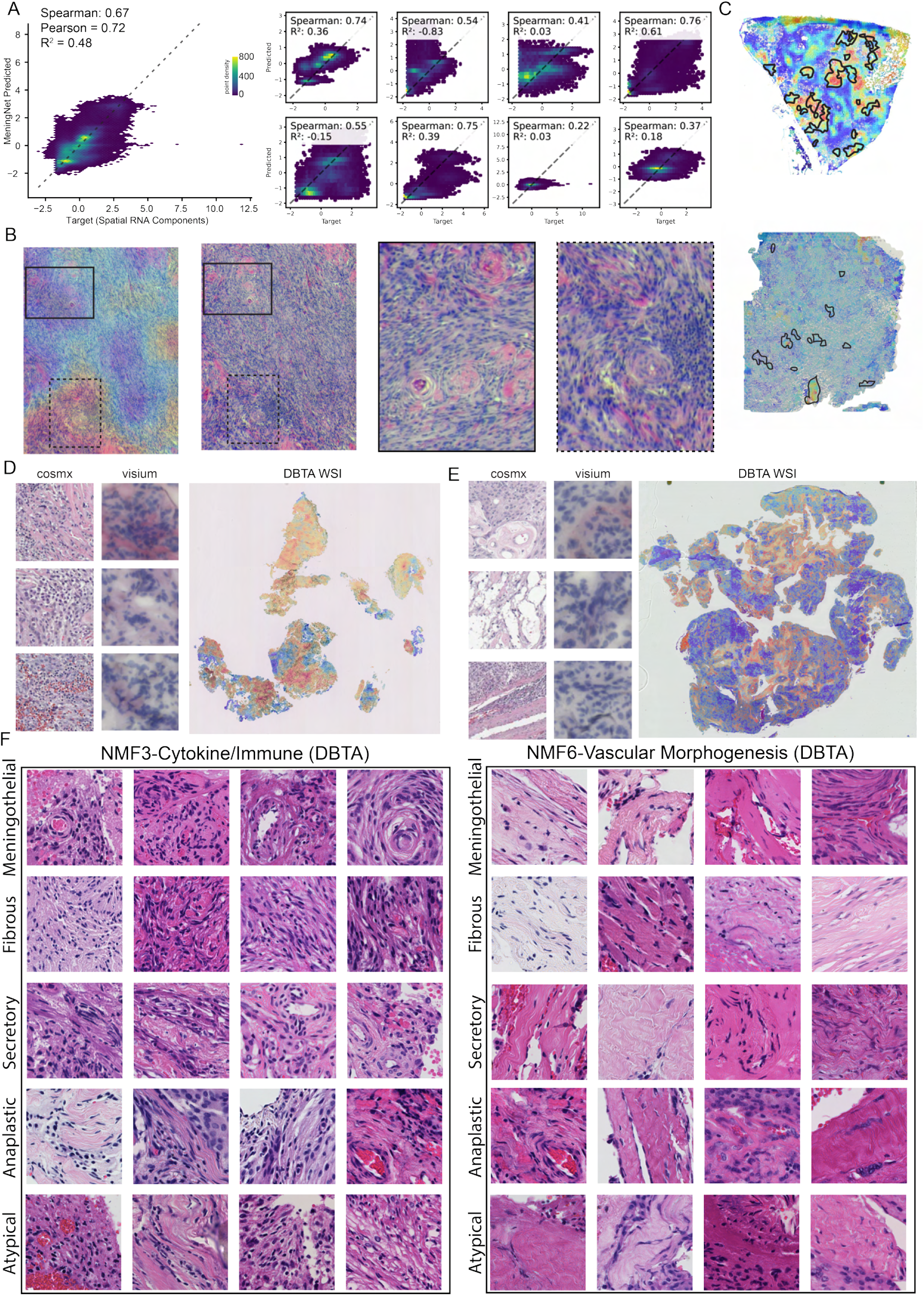
Morphological Representations of SMPs. A. Correlation density plots summarizing aggregated model performance (left) and per-component performance breakdown (right). **B.** Inferred SMP map (left) for RNA-NMF3 (cytokine/immune) in a representative sample from the internal replication cohort. demonstrating two spatially distinct regions of immune infiltration. Boxes indicate higher-resolution regions shown at right, differentiated by solid and dashed outlines. **C.** Inferred SMP map of RNA-NMF2 (oxidative phosphorylation) with whorl–lobule (WL) score contours overlaid on Yale-NS-0848 (top) and Yale-NS-0856 (bottom) samples. **D.** Representative patches from CosMx (left) and Visium (middle), alongside DBTA whole-slide inference maps (right) for RNA-NMF3 (cytokine/immune). **E.** DBTA whole-slide inference map for RNA-NMF6 (vascular morphogenesis). **F.** Representative top-scoring image patches for RNA-NMF3 (cytokine/immune) (left) and RNA-NMF6 (vascular morphogenesis) (right) across histologic subtypes.

**Figure S11.**
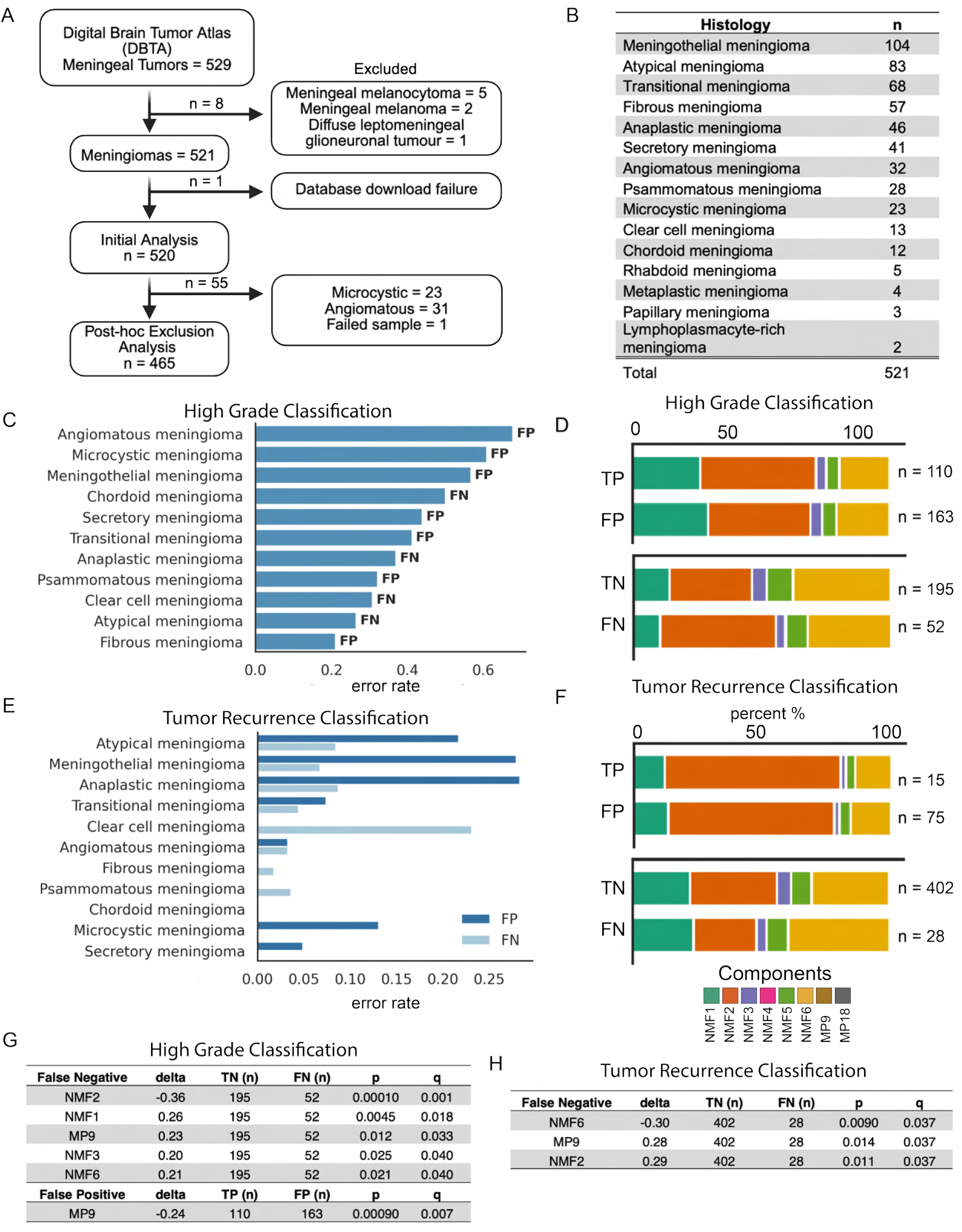
Digital Brain Tumor Atlas Validation Cohort. **A.** Cohort curation and exclusion workflow. From 529 meningeal tumors in DBTA, non-meningioma entities and download failures were excluded, resulting in n = 521 meningiomas. This was followed by post-hoc exclusion of selected histologic subtypes, yielding the final analysis cohort of n = 465. **B.** Distribution of meningioma histologic subtypes available from the Digital Brain Tumor Atlas (DBTA) whole-slide H&E cohort (n = 521 after initial filtering). **C.** Post-hoc analysis of grade classification errors stratified by histologic subtype, showing error rate separated into false positive (FP; low-grade predicted as high-grade) and false negative (FN; high-grade predicted as low-grade) proportions. **D.** Distribution of dominant inferred spatial molecular components (SMPs) among true positives (TP), false positives (FP), true negatives (TN), and false negatives (FN) for grade classification. **E.** Post-hoc analysis of recurrence classification errors stratified by histologic subtype, separated into FP and FN proportions. **F.** Distribution of dominant inferred SMPs among TP, FP, TN, and FN for recurrence prediction. **G.** Component-level enrichment analysis identifying SMPs significantly associated with false negative and false positive classifications for grade (delta indicates difference in component usage between error class and correctly classified tumors; p and q values shown). **H.** Component-level enrichment analysis identifying SMPs significantly associated with false negative classifications for recurrence prediction.

## Supplemental Files

Supplementary File 1

Supplementary File 2

Supplementary File 3

Supplementary File 4

Supplementary File 5

Supplementary File 6

Supplementary File 7

Supplementary File 8

Supplementary File 9

